# Outer membrane stiffness gates multidrug tolerance

**DOI:** 10.64898/2026.08.24.746625

**Authors:** Nan Xiao, Bo Zheng, Jia-Feng Liu

## Abstract

Bactericidal antibiotics initiate killing through class-specific target damage, yet the cellular properties that determine whether antibiotic-induced damage remains reversible or progresses to irreversible death remains unclear. Here, using kanamycin-centered evolution in *Escherichia coli*, we identified a multidrug-tolerant mutant that exhibits increased survival across aminoglycosides, β-lactams, quinolones and polymyxins without altered minimum inhibitory concentrations. We demonstrate that mechanistically distinct antibiotics converge on outer membrane destabilization, revealing a shared downstream vulnerability during killing. Elevated outer membrane stiffness limits antibiotic-induced envelope destabilization, thereby gating multidrug tolerance. Orthogonal chemical and physical perturbations further established a quantitative relationship between outer membrane stiffness and antibiotic survival across drug classes. Our findings reveal outer membrane stiffness as a previously unrecognized physical basis of multidrug tolerance and suggest that modulating bacterial envelope mechanics may provide new opportunities for antimicrobial intervention.

## Introduction

How bactericidal antibiotics kill bacteria remains a central question in microbiology. Antibiotic lethality is a dynamic process that begins with primary target engagement and disruption of essential cellular functions, propagates through downstream damage and culminates in irreversible death ^1,2^. Although drug–target interactions and their immediate consequences have been extensively characterized, the processes that determine whether antibiotic-induced damage remains reparable or progresses to irreversible death remain poorly understood ^3,4^. The continued spread of multidrug resistance and the lack of new antibiotics further heighten the need to identify cellular vulnerabilities that can be exploited to improve bacterial killing ^5,6^. This need is particularly acute in Gram-negative pathogens, whose outer membrane (OM) constitutes an intrinsic barrier to many antibiotics and thereby restricts the repertoire of effective antibacterial agents ^7,8^. Therefore, defining the cellular determinants that govern the transition from antibiotic-induced damage to irreversible death in Gram-negative bacteria could reveal strategies to potentiate existing antibiotics and identify new antibacterial targets.

Antibiotic tolerance provides an informative framework for dissecting this transition. Unlike antibiotic resistance, which increases the minimum inhibitory concentration (MIC) and permits bacterial growth at drug concentrations that inhibit susceptible cells ^9,10^, tolerance prolongs the time required to kill a susceptible population even at high antibiotic concentrations ^11,12^. By uncoupling killing kinetics from growth inhibition, tolerance enables the identification of cellular processes that determine survival after antibiotic-induced damage. Tolerance to individual antibiotics can be mediated by diverse physiological processes, including growth arrest, stress-response activation, metabolic remodeling and damage repair ^13–16^. Nevertheless, what ultimately prevents antibiotic-induced damage from progressing to irreversible death remains unclear. Cross-tolerance between antibiotics acting on unrelated primary targets has long been observed ^17–19^. Previous studies have shown that such cross-protection can arise through both antibiotic-specific processes operating in parallel and shared protective mechanisms acting across antibiotic classes ^20–22^. Such shared mechanisms raise the possibility that a common cellular determinant may confer multidrug tolerance by limiting the progression of distinct forms of antibiotic-induced damage to irreversible death. However, the cellular properties that enable such broad protection in Gram-negative bacteria remain to be identified.

Here, using a kanamycin-centered evolution strategy in *Escherichia coli*, we identified a multidrug-tolerant mutant and uncovered its cellular basis. Single-cell dual-fluorescence imaging revealed that mechanistically distinct bactericidal antibiotics converge on OM destabilization before cytoplasmic failure, whereas multidrug-tolerant cells preserve OM stability during killing. Indentation-depth-resolved atomic force microscopy (AFM) further demonstrated a preferential increase in OM stiffness in multidrug-tolerant cells. By independently tuning OM mechanics through chemical and physical perturbations, we established a quantitative relationship between OM stiffness and antibiotic survival across drug classes. These findings identify OM stiffness as a physical determinant of multidrug tolerance and suggest that bacterial envelope mechanics may represent a potential target for new antibacterial strategies.

## Results

### Kanamycin persisters exhibit broad protection across antibiotic classes

To investigate whether tolerance can extend across antibiotics with distinct primary mechanisms of action, we performed reciprocal persister cross-challenge assays in *Escherichia coli* (*E. coli*) KLY using three representative antibiotics: ampicillin (β-lactam), norfloxacin (quinolone) and kanamycin (aminoglycoside), which primarily inhibit cell-wall synthesis, DNA replication and translation, respectively ^1^. Persisters surviving an initial exposure to ampicillin, norfloxacin or kanamycin were washed and then separately rechallenged with each of the three antibiotics (Extended Data Fig. 1a). Compared with the untreated population, ampicillin and norfloxacin persisters showed reduced killing upon rechallenge with either ampicillin or norfloxacin but remained efficiently killed by kanamycin (Fig. 1a). By contrast, kanamycin persisters showed reduced killing not only upon kanamycin rechallenge but also upon subsequent exposure to ampicillin or norfloxacin (Fig. 1a). These asymmetric survival profiles suggest that kanamycin treatment preferentially selects persisters with broader protection across antibiotic classes.

**Fig. 1.**
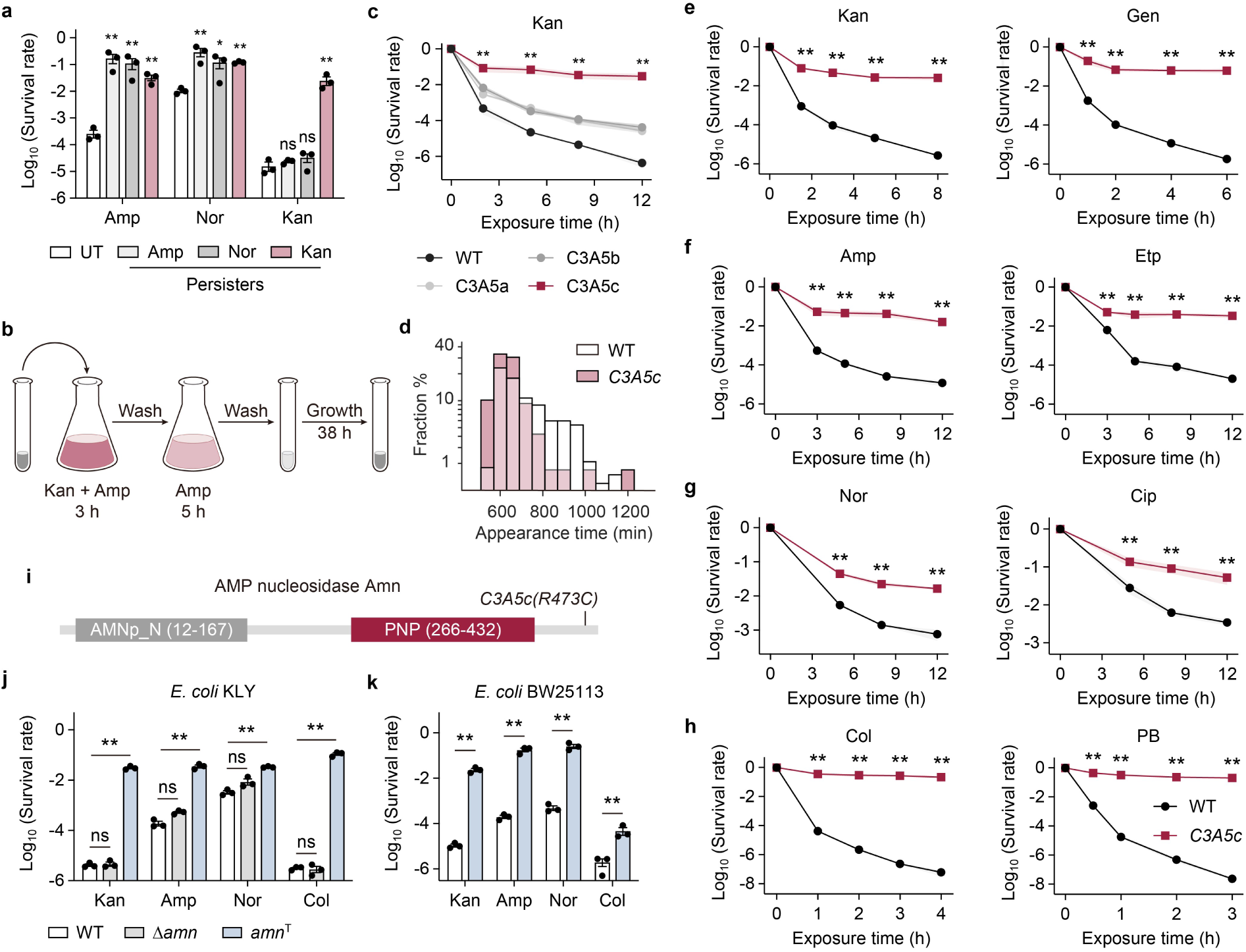
Kanamycin-centered evolution uncovers genetically encoded multidrug tolerance. **a**, Cross-challenge survival of the untreated (UT) *Escherichia coli* KLY wild-type cells and persisters isolated after primary treatment with ampicillin (Amp; 5 h), norfloxacin (Nor; 5 h) or kanamycin (Kan; 5 h), followed by challenge with the indicated antibiotics. **b**, Schematic of the kanamycin-centered evolution. In each cycle, parallel cultures were diluted 100-fold into fresh LB, and treated with kanamycin and ampicillin for 3 h, washed and exposed to ampicillin for an additional 5 h. After antibiotic removal, survivors were resuspended in fresh LB and grown for 38 h before the next cycle. **c**, Kanamycin time-kill curves of the ancestral wild type (WT) and independently evolved clones *C3A5a*, *C3A5b* and *C3A5c*. **d**, Colony appearance-time distributions of wild-type (*n* = 275 colonies) and *C3A5c* (*n* = 240 colonies) cells under antibiotic-free conditions, determined by ScanLag. **e**–**h**, Time-kill curves comparing wild-type and *C3A5c* cells under treatment with aminoglycosides (kanamycin; gentamicin, Gen; **e**), β-lactams (ampicillin; ertapenem, Etp; **f**), quinolones (norfloxacin; ciprofloxacin, Cip; **g**), or polymyxins (colistin, Col; polymyxin B, PB; **h**). **i**, Domains of AMP nucleosidase Amn. The R473C substitution in *C3A5c* is indicated. AMN_N, Amn N-terminal domain; PNP, purine nucleoside phosphorylase. **j**, **k**, Survival of wild-type, *amn*-deletion (Δ*amn*) and reconstructed *amn* R473C (*amn*^T^) strains in *E. coli* KLY (**j**) and BW25113 (**k**) backgrounds following treatment with kanamycin (5 h), ampicillin (5 h), norfloxacin (8 h) or colistin (2 h). Antibiotic treatments were performed with kanamycin (120 µg/ml), gentamicin (50 µg/ml), ampicillin (120 µg/ml), ertapenem (2 µg/ml), norfloxacin (8 µg/ml), ciprofloxacin (8 µg/ml), colistin (10 µg/ml) or polymyxin B (10 µg/ml). Survival data are mean ± s.e.m. from three biological replicates. Statistical analysis was performed using two-way ANOVA with Bonferroni correction. ns, not significant; \**P* < 0.05; \*\**P* < 0.01.

### Kanamycin-centered evolution yields multidrug tolerance

To determine whether the broad protection could be genetically stabilized, we adapted an evolution strategy previously shown to enrich antibiotic-tolerant mutants ^18,23^. We first subjected KLY to cyclic kanamycin exposure at 120 µg ml^-1^, corresponding to 20× the minimum inhibitory concentration (MIC) (Extended Data Fig. 1b). This scheme rapidly enriched kanamycin-resistant cells within two to three cycles, producing a pronounced increase in population survival (Extended Data Fig. 1c, d). However, these resistant cells remained susceptible to ampicillin and showed no survival advantage during ampicillin treatment (Extended Data Fig. 1c, d). We therefore reasoned that ampicillin could serve as a counterselection pressure to restrict the expansion of kanamycin-resistant cells. Accordingly, we modified the evolution scheme by imposing a sequential ampicillin challenge after combined kanamycin and ampicillin exposure (Fig. 1b), thereby constraining resistance-driven outgrowth while allowing the enrichment of tolerant survivors.

After five cycles, all three independently evolved populations, *C3A5a*, *C3A5b* and *C3A5c*, showed increased survival during kanamycin treatment (Extended Data Fig. 2a). Clones isolated from these evolved populations retained the same kanamycin MIC as the wild type (Extended Data Fig. 2b), indicating that their enhanced survival was not attributable to kanamycin resistance. We therefore examined their time-kill dynamics to determine whether the increased survival reflected kanamycin tolerance ^11^. The wild type, *C3A5a* and *C3A5b* reached 99% killing within 2 h of kanamycin treatment, whereas *C3A5c* required more than sixfold longer to reach comparable killing (Fig. 1c). This prolonged killing duration was not accompanied by detectable shifts in colony appearance-time distribution or growth rate under drug-free conditions (Fig.1d and Extended Data Fig. 2c), distinguishing *C3A5c* from previously described tolerance associated with pre-existing lag extension or constitutively slow growth. Thus, these results establish *C3A5c* as a bona fide kanamycin-tolerant strain.

To determine whether *C3A5c* recapitulated the broad cross-class protection observed in kanamycin persisters, we measured MICs and time-kill dynamics across aminoglycosides, β-lactams, quinolones and polymyxins. MICs for all tested antibiotics were indistinguishable between *C3A5c* and the wild type (Extended Data Fig. 2d–g and Supplementary Table 1), excluding resistance as the basis of enhanced survival. During kanamycin and gentamicin treatment, *C3A5c* exhibited delayed killing and increased population survival (Fig. 1e), indicating increased tolerance across aminoglycosides. Increased tolerance was also observed during treatment with the β-lactams ampicillin and ertapenem and the quinolones norfloxacin and ciprofloxacin (Fig. 1f, g). Notably, *C3A5c* further exhibited increased tolerance to the polymyxins colistin and polymyxin B, last-resort antibiotics used against multidrug-resistant Gram-negative pathogens (Fig. 1h). Collectively, these data establish multidrug tolerance in *C3A5c* across four mechanistically distinct classes of bactericidal antibiotics.

### *amn*^T^-mediated multidrug tolerance is not explained by loss of canonical Amn catalysis

Whole-genome sequencing of *C3A5c* identified a single-nucleotide variant in *amn* (hereafter *amn*^T^) that resulted in an R473C substitution in the encoded AMP nucleosidase Amn (Fig. 1i). Reconstruction of the *amn*^T^ allele in the ancestral KLY background reproduced the multidrug-tolerant phenotype of *C3A5c* (hereafter the *amn*^T^ mutant) (Fig. 1j and Extended Data Fig. 3a). Consistently, the *E. coli* BW25113 *amn*^T^ strain also showed increased antibiotic tolerance (Fig. 1k and Extended Data Fig. 3b), demonstrating that *amn*^T^ allele is sufficient to confer multidrug tolerance across distinct *E. coli* backgrounds.

Because Amn catalyzes the hydrolysis of AMP into adenine and ribose 5-phosphate ^24^, we asked whether the tolerant phenotype resulted from altered canonical Amn activity. In vitro enzymatic assays showed that Amn^T^ had impaired AMP nucleosidase activity relative to wild-type Amn (Extended Data Fig. 3c, d). However, deletion of *amn* (Δ*amn*) did not increase antibiotic tolerance (Fig. 1j and Extended Data Fig. 3a). Moreover, IPTG-induced expression of the catalytically impaired Amn D428A variant in the Δ*amn* background failed to reproduce the tolerant phenotype (Extended Data Fig. 3e, f). Thus, although R473C substitution reduces Amn catalytic activity, loss of canonical Amn catalysis is not sufficient to confer multidrug tolerance.

### Multidrug-tolerant cells are protected from shared outer-membrane destabilization during antibiotic killing

The multidrug tolerance conferred by *amn*^T^ prompted us to ask whether it reflects protection against a shared cellular vulnerability exposed by mechanistically distinct bactericidal antibiotics. To examine this possibility, we first assessed representative cellular processes previously linked to antibiotic killing, including reactive oxygen species (ROS) accumulation, translational inhibition, DNA-damage signaling and membrane permeabilization ^1,2,4^. ROS production, assessed by dihydroethidium staining, increased after antibiotic treatment to comparable levels in wild-type and *amn*^T^ cells (Extended Data Fig. 4a). Similarly, de novo protein synthesis, monitored by cytoplasmic YFP accumulation after transfer to fresh LB medium, was completely inhibited by kanamycin in both strains (Extended Data Fig. 4b). Norfloxacin-induced SOS activation, measured by P*sulA*-mCherry fluorescence, was also comparable between wild-type and *amn*^T^ cells (Extended Data Fig. 4c). By contrast, outer-membrane (OM) and inner-membrane (IM) permeability, measured by N-phenyl-1-naphthylamine (NPN) and propidium iodide (PI) uptake, respectively, increased markedly in wild-type cells during ampicillin treatment, whereas both permeability defects were attenuated in *amn*^T^ cells (Extended Data Fig. 5a).

To examine this envelope protection at the single-cell level, we monitored wild-type and *amn*^T^ cells during ampicillin exposure using a dual-fluorescence reporter system comprising periplasmic mCherry and cytoplasmic YFP. Consistent with β-lactam-induced cell-wall damage and turgor-driven envelope stress ^25,26^, wild-type cells underwent progressive swelling, with steady increases in both cell width and length (Fig. 2a, b and Supplementary Video 1). This swelling was followed by YFP-labeled cytoplasmic bulging and culminated in either lysis (Fig. 2c, d) or severe irregular deformation (Fig. 2e, f and Extended Data Fig. 5b). Crucially, in both lytic and severely deformed cells, dual-reporter imaging revealed that focal redistribution of periplasmic mCherry consistently preceded cytoplasmic bulging (Fig. 2d, f). Moreover, cytoplasmic bulging appeared later in severely deformed cells than in cells that proceeded to lysis, accompanied by a corresponding delay in focal periplasmic mCherry redistribution (Extended Data Fig. 5c). This temporal ordering suggests a damage sequence in which local OM destabilization precedes overt cytoplasmic deformation during ampicillin treatment. Consistently, approximately 90% of wild-type cells progressed to lysis or severe deformation, whereas more than 70% of *amn*^T^ cells showed no detectable swelling and maintained stable periplasmic mCherry distribution throughout 3 h of ampicillin exposure (Fig. 2a, b, g, Extended Data Fig. 5d and Supplementary Video 2), indicating markedly preserved OM stability in *amn*^T^ cells.

**Fig. 2.**
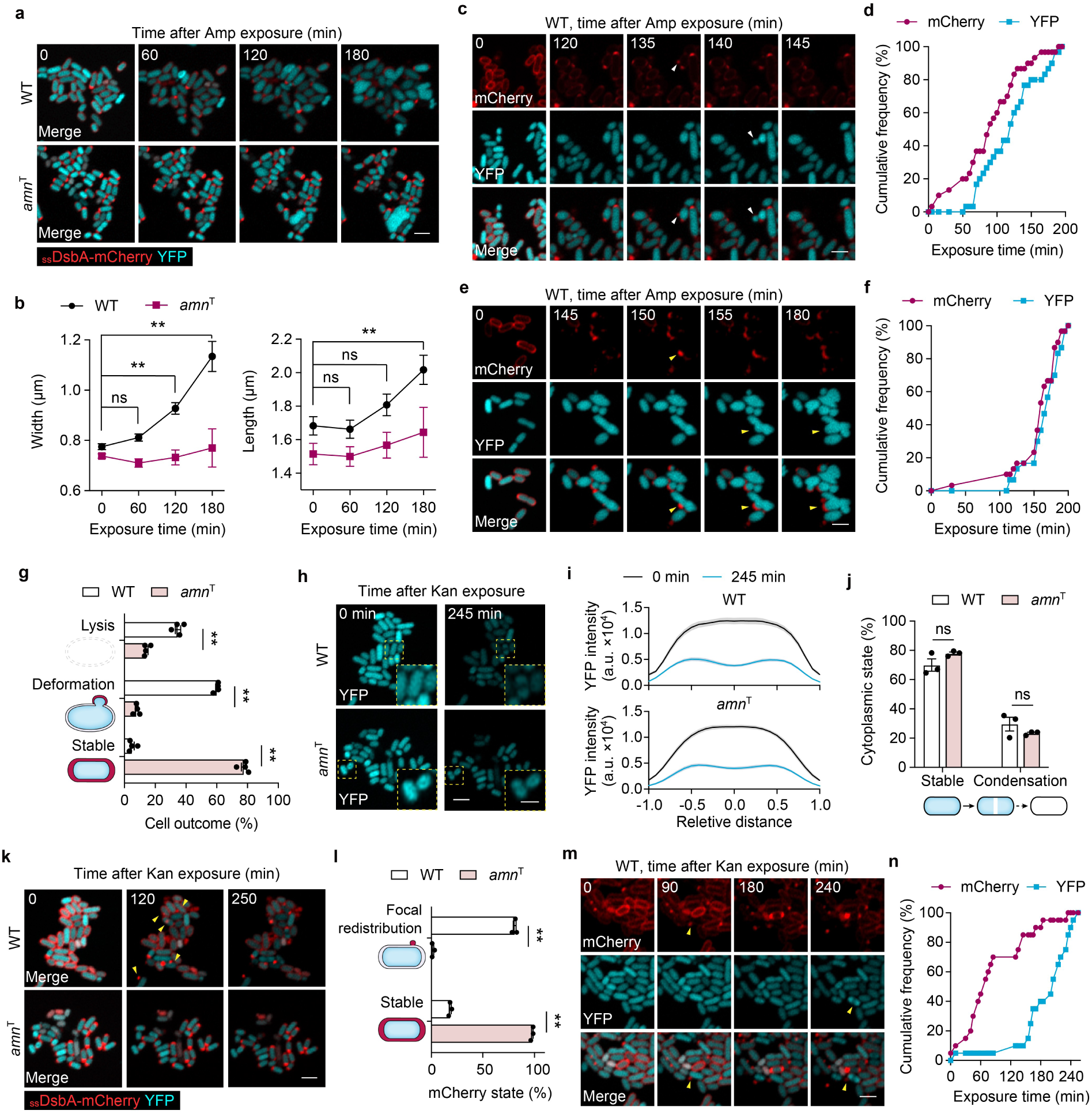
Mechanistically distinct bactericidal antibiotics expose a shared downstream outer-membrane vulnerability. **a**, Representative time-lapse fluorescence images of wild-type and *amn*^T^ cells expressing periplasmic _ss_DsbA–mCherry (red) and cytoplasmic yellow fluorescent protein (YFP; cyan) during ampicillin exposure. **b**, Cell width and length of wild-type (*n* = 34 cells) and *amn*^T^ (*n* = 25 cells) cells during ampicillin exposure. **c**–**f**, Temporal relationship between periplasmic and cytoplasmic alterations during ampicillin exposure in wild-type cells. **c**, **e**, Representative single-cell time-lapse sequences showing changes in periplasmic mCherry and cytoplasmic YFP signals in cells undergoing lysis (**c**) or non-lytic deformation (**e**). Arrowheads indicate the corresponding fluorescence or morphological transitions. **d**, **f**, Cumulative frequencies of the onset times of the corresponding mCherry and YFP events shown in **c** and **e**, respectively (*n* = 20 cells per group). **g**, Frequencies of cell outcomes after 180 min of ampicillin exposure, classified as lysis, severe deformation or maintenance of a stable rod-like morphology (*n* = 4 trials; 90–210 cells per group). **h**, **i**, Representative YFP fluorescence images (**h**) and longitudinal YFP fluorescence-intensity profiles (**i**; *n* = 20 cells per group) of wild-type and *amn*^T^ cells before and after 245 min of kanamycin exposure. Insets show enlarged views of the boxed regions. Cell length was normalized from −1 to 1. **j**, Frequencies of cells maintaining a stable cytoplasmic YFP distribution or undergoing YFP condensation after 245 min of kanamycin exposure (*n* = 3 trials; 60–110 cells per group). **k**, Representative time-lapse fluorescence images of wild-type and *amn*^T^ cells during kanamycin exposure, showing periplasmic _ss_DsbA–mCherry (red) and cytoplasmic YFP (cyan). Arrowheads indicate focal redistribution of periplasmic mCherry. **l**, Frequencies of cells showing focal redistribution of periplasmic mCherry or maintaining a stable periplasmic fluorescence pattern after 245 min of kanamycin exposure (*n* = 3 trials; 60–110 cells per group). **m**, Representative time-lapse sequence of a wild-type cell showing focal redistribution of periplasmic mCherry preceding cytoplasmic YFP alteration during kanamycin exposure. Arrowheads indicate the corresponding events. **n**, Cumulative frequencies of the onset times of periplasmic mCherry redistribution and cytoplasmic YFP alteration in wild-type cells during kanamycin exposure (*n* = 20 cells). Ampicillin and kanamycin were used at 120 µg/ml. Data are mean ± s.e.m. Statistical analysis was performed using two-way ANOVA with Bonferroni correction; ns, not significant; \**P* < 0.05; \*\**P* < 0.01. Scale bars, 2 µm (a, c, e, h, k, m) and 1 µm (h, insets).

We next asked whether OM destabilization also occurs during aminoglycoside killing, which, beyond its primary effect on translation, has been associated with membrane damage and cytoplasmic condensation ^27–29^. During kanamycin treatment, wild-type and *amn*^T^ cells showed comparable frequencies of cytoplasmic condensation (Fig. 2h–j and Extended Data Fig. 6a), indicating that this cytoplasmic response does not account for their differential survival. Although bulk NPN and PI uptake remained minimal during kanamycin treatment (Extended Data Fig. 6b), time-lapse imaging revealed focal redistribution of periplasmic mCherry in approximately 80% of wild-type cells (Fig. 2k, l and Supplementary Video 3). Moreover, among wild-type cells undergoing cytoplasmic condensation, focal mCherry redistribution consistently preceded YFP-labeled condensation (Fig. 2m, n), indicating that local OM destabilization occurs before detectable cytoplasmic condensation. By contrast, focal mCherry redistribution was observed in only approximately 2% of *amn*^T^ cells, with more than 90% maintaining a stable periplasmic mCherry distribution throughout 5 h of kanamycin exposure (Fig. 2k, l, Extended Data Fig. 6c and Supplementary Video 4). Consistently, scanning electron microscopy revealed surface protrusions in kanamycin-treated wild-type cells but rarely in *amn*^T^ cells (Extended Data Fig. 6d). These observations reveal a previously underappreciated local OM-destabilization event during kanamycin killing that is strongly attenuated in *amn*^T^ cells.

Collectively, the selective attenuation of OM destabilization in *amn*^T^ cells, rather than of antibiotic-specific intracellular responses, supports OM destabilization as a shared downstream vulnerability in the progression from antibiotic-induced damage to irreversible death.

### Elevated outer membrane stiffness limits envelope destabilization and gates multidrug tolerance

Recent studies have shown that the OM bears substantial mechanical loads in Gram-negative bacteria and that its stiffness limits envelope deformation and lysis during mechanical perturbation ^30,31^. To determine whether the *amn*^T^ mutation alters OM mechanics, we directly probed the stiffness of intact cells using atomic force microscopy (AFM). Stiffness was first estimated within the initial ∼10 nm indentation range (Fig. 3a and Extended Data Fig. 7a, b), a shallow depth expected to be enriched for OM-associated mechanical contributions based on envelope geometry ^32^ (Extended Data Fig. 7c, d). Within this range, *amn*^T^ cells exhibited significantly higher stiffness than wild-type cells (Fig. 3b). Notably, this stiffness difference was most pronounced near the cell surface and progressively diminished with deeper indentation, with no appreciable difference remaining by approximately 20 nm (Fig. 3c, d). Consistent with this depth-dependent interpretation, deletion of *lpp* (Δ*lpp*), which encodes the major lipoprotein tethering the OM to peptidoglycan and contributes to envelope mechanics ^30,33^, had little effect at shallow indentation (∼10 nm) but reduced stiffness at 15–20 nm (Extended Data Fig. 7e). Taken together, these measurements support a preferential increase in OM-associated stiffness in *amn*^T^ cells rather than a uniform increase in whole-envelope rigidity.

**Fig. 3.**
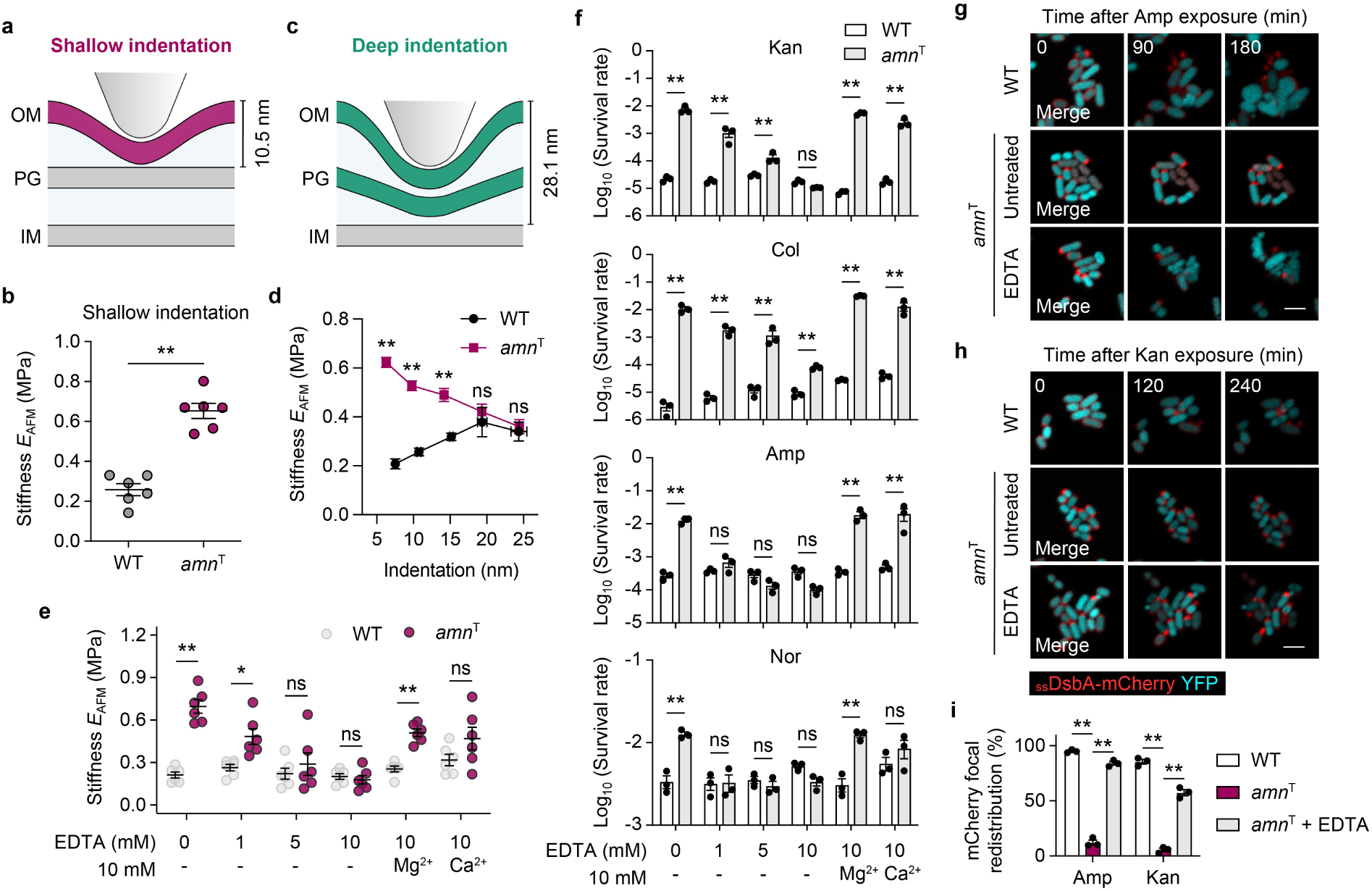
Outer-membrane stiffness gates multidrug tolerance. **a**, Schematic of stiffness measurements under shallow atomic force microscopy (AFM) indentation. OM, outer membrane; PG, peptidoglycan; IM, inner membrane. **b**, Stiffness *E*_AFM_ of wild-type and *amn*^T^ cells measured under shallow indentation. Each point represents the mean stiffness of an individual cell (*n* = 6 cells per group). **c**, Schematic of stiffness measurements under deeper AFM indentation. **d**, Stiffness of wild-type and *amn*^T^ cells measured at indicated indentation depths (*n* = 6 cells per group). **e**, Stiffness of wild-type and *amn*^T^ cells after 1 h of treatment with the indicated concentrations of EDTA, with or without 10 mM MgCl_2_ or CaCl_2_ (*n* = 6 cells per group). **f**, Survival of wild-type and *amn*^T^ cells following the indicated EDTA treatments, with or without 10 mM MgCl_2_ or CaCl_2_, followed by challenge with kanamycin (120 µg/ml, 5 h), colistin (10 µg/ml, 2 h), ampicillin (120 µg/ml, 5 h) or norfloxacin (8 µg/ml, 8 h). *n* = 3 biological replicates per group. **g**, **h**, Representative time-lapse fluorescence images of wild-type, untreated *amn*^T^ and 10 mM EDTA-treated *amn*^T^ cells during ampicillin (**g**) or kanamycin (**h**) exposure. Periplasmic _ss_DsbA–mCherry is shown in red and cytoplasmic YFP in cyan. Scale bars, 2 µm. **i**, Frequencies of cells exhibiting focal redistribution of periplasmic mCherry during ampicillin or kanamycin exposure in wild-type, untreated *amn*^T^ and 10 mM EDTA-treated *amn*^T^ cells (*n* = 3 trials; 60– 90 cells per group). Data are mean ± s.e.m. Statistical analysis was performed using a two-sided unpaired Student’s t-test (**b**) or two-way ANOVA with Bonferroni correction (**e**, **f**, **i**); ns, not significant; \**P* < 0.05; \*\**P* < 0.01.

To determine whether elevated OM stiffness functionally supports multidrug tolerance, we transiently perturbed OM stabilization with EDTA, which chelates the divalent cations that electrostatically bridge adjacent LPS molecules ^34^. Short-term EDTA treatment resulted in reduced OM stiffness of *amn*^T^ cells in a concentration-dependent manner, whereas supplementation with excess Mg^2+^ or Ca^2+^ prevented this reduction (Fig. 3e and Extended Data Fig. 8a). Following this transient chelation pulse, cells were washed and transferred to fresh medium before antibiotic challenge to minimize direct effects of EDTA on antibiotic activity. EDTA treatment alone did not measurably alter cell viability or lag time (Extended Data Fig. 8b, c). Nevertheless, increasing EDTA concentrations progressively eroded the survival advantage of *amn*^T^ cells across multiple antibiotics without altering MICs (Fig. 3f and Extended Data Fig. 8d), indicating a loss of tolerance rather than a change in resistance. Conversely, Mg²⁺ or Ca²⁺ repletion restored the survival advantage of *amn*^T^ cells (Fig. 3f).

To further link elevated OM stiffness to protection against antibiotic-induced OM destabilization, we examined EDTA-treated *amn*^T^ cells by time-lapse microscopy. Consistent with the antibiotic-induced OM destabilization observed in wild-type cells, EDTA treatment markedly increased the fraction of *amn*^T^ cells exhibiting focal periplasmic mCherry redistribution during both ampicillin and kanamycin exposure (Fig. 3g–i and Supplementary Video 5, 6). Collectively, the coordinated changes in OM stiffness, envelope stability and survival support a functional role for elevated OM stiffness in *amn*^T^-mediated multidrug tolerance.

### Orthogonal perturbations reveal a quantitative stiffness–survival relationship

Having established that elevated OM stiffness contributes to multidrug tolerance, we next asked whether an orthogonal physical perturbation could tune OM mechanics and thereby modulate antibiotic survival. Given that osmotic shifts can rapidly alter cell hydration, turgor and envelope prestress ^35,36^, we transiently exposed stationary-phase cells to a graded series of sorbitol concentrations for 1 h, followed by sorbitol washout before AFM measurements or antibiotic challenge (Fig. 4a). Transient exposure to 0.3 M sorbitol increased the width of wild-type and *amn*^T^ cells (Fig. 4b, c and Extended Data Fig. 9a), consistent with osmotic rehydration and envelope stretching. This treatment alone did not affect cell viability or antibiotic susceptibility (Extended Data Fig. 9b, c), but markedly reduced the elevated OM stiffness of *amn*^T^ cells and largely abolished their tolerance advantage (Fig. 4d, e and Extended Data Fig. 9d). Increasing sorbitol concentrations progressively restored both stiffness and survival, with both approaching untreated *amn*^T^ levels at 1.5 M (Fig. 4d, e). Across the osmotic series, survival during treatment with multiple antibiotics was strongly correlated with OM stiffness (Extended Data Fig. 9e), establishing a quantitative stiffness–survival coupling under an independent physical perturbation.

**Fig. 4.**
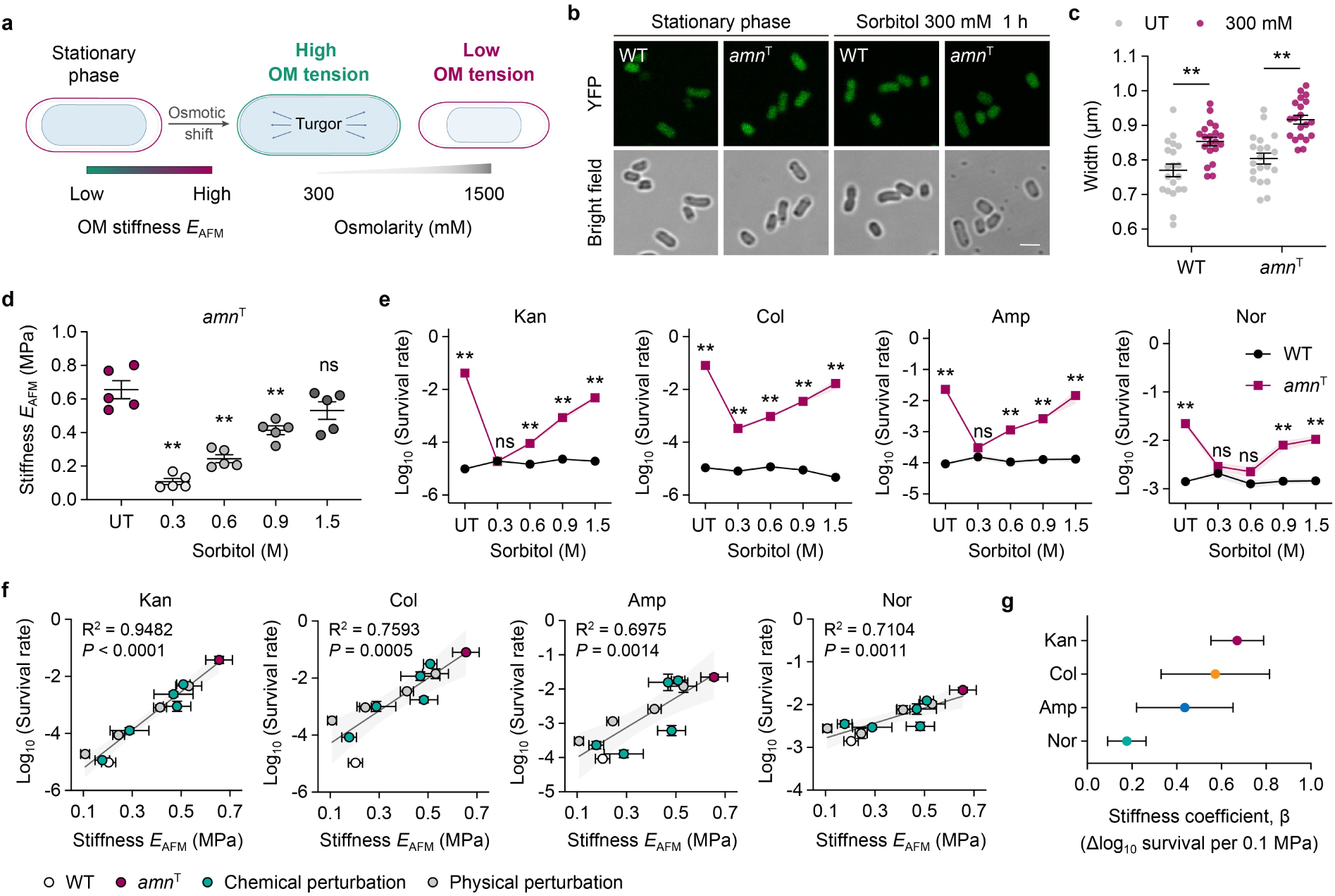
Antibiotic survival quantitatively scales with outer-membrane stiffness across orthogonal perturbations. **a**, Schematic of osmotic modulation of outer-membrane mechanics. Changes in external osmolarity may alter cell turgor and outer-membrane tension, thereby modulating outer-membrane stiffness. **b**, Representative cytoplasmic YFP fluorescence (green) and bright-field images of stationary-phase wild-type and *amn*^T^ cells before and after treatment with 300 mM sorbitol for 1 h. Scale bar, 2 µm. **c**, Cell width of wild-type and *amn*^T^ cells before (UT) and after treatment with 300 mM sorbitol for 1 h (*n* = 20 cells per group). **d**, Stiffness of *amn*^T^ cells before and after treatment with the indicated sorbitol concentrations for 1 h (*n* = 5 cells per group). **e**, Survival of wild-type and *amn*^T^ cells following treatment with the indicated sorbitol concentrations for 1 h and subsequent challenge with kanamycin (120 µg/ml, 5 h), colistin (10 µg/ml, 2 h), ampicillin (120 µg/ml, 5 h), or norfloxacin (8 µg/ml, 8 h). *n* = 3 biological replicates per group. **f**, Relationship between stiffness and antibiotic survival across genetic, chemical and physical perturbations for kanamycin, colistin, ampicillin and norfloxacin. Solid lines indicate linear regression fits, with R^2^ and *P* values shown. **g**, Stiffness coefficients (β) estimated from linear regression models in **f** for each antibiotic. Coefficients represent the change in log_10_ survival associated with a 0.1-MPa increase in stiffness. Error bars indicate 95% confidence intervals. Data are mean ± s.e.m. Statistical analysis was performed using one-way ANOVA (**d**) or two-way ANOVA (**c** and **e**) with Bonferroni correction, and linear regression (**f**); ns, not significant; \*\**P* < 0.01.

To determine whether this quantitative relationship was conserved across distinct modes of envelope perturbation, we integrated stiffness and survival measurements from osmotic preconditioning with those from EDTA treatment and divalent-cation supplementation. For each of the four antibiotics, chemical and physical perturbations converged on similar positive linear relationships between OM stiffness and log_10_ survival (Fig. 4f and Extended Data Fig. 9f), indicating that the stiffness–survival coupling was preserved across independent modes of perturbation. Nevertheless, the slopes of these relationships differed among antibiotics, yielding drug-specific stiffness coefficients, β, defined as the fitted change in log_10_ survival associated with a 0.1-MPa increase in OM stiffness (Fig. 4g and Extended Data Fig. 9f). Thus, although increased OM stiffness consistently favored survival, the magnitude of this benefit was antibiotic dependent. Taken together, these results establish OM stiffness as a common quantitative determinant of multidrug tolerance across orthogonal perturbations.

## Discussion

In this work, we identify OM stiffness as a physical determinant of multidrug tolerance in *Escherichia coli*. Our findings show that multidrug-tolerant cells do not survive by suppressing canonical intracellular damage, but by maintaining a mechanically stabilized OM that limits antibiotic-induced OM destabilization during exposure to mechanistically distinct antibiotics. By independently modulating OM mechanics, we further establish that OM stiffness quantitatively influences antibiotic survival across antibiotic classes. Together, our findings reveal bacterial OM mechanics as a previously underappreciated determinant of how antibiotic-induced damage progresses to irreversible death.

Despite a growing understanding of antibiotic-induced cellular damage, the factors that determine whether such damage remains reversible or progresses to irreversible death remain poorly understood ^2^. An influential model proposed that bactericidal antibiotics converge on ROS-mediated killing, although the generality and causal contribution of this pathway remain controversial ^37–40^. In our study, multidrug-tolerant cells retained antibiotic-induced ROS production, translational inhibition and DNA-damage responses at levels comparable to wild-type cells, indicating that multidrug tolerance arises independently of changes in these canonical intracellular damage responses. Instead, multidrug-tolerant cells were distinguished by their ability to preserve OM stability during antibiotic challenge. This concept parallels observations in eukaryotic lytic cell death, where NINJ1 controls plasma-membrane rupture and defines a mechanical threshold for membrane failure ^41^. In bacteria, however, it remains unresolved how antibiotic-induced damage is translated into OM destabilization. Antibiotics may directly perturb the OM or generate cellular stresses that mechanically overload the envelope beyond its load-bearing capacity. For β-lactams, inhibition of PG synthesis compromises cell-wall integrity and exposes the envelope to turgor-driven deformation and lysis ^25,26^, while disruption of Lpp-mediated PG–OM attachment can further destabilize the envelope ^42^. Aminoglycosides, although primarily targeting translation, have also been associated with envelope perturbation through direct interactions with the OM and mistranslation-induced membrane defects that impair membrane integrity and promote further antibiotic uptake ^27,28,43^. Recent single-cell studies further linked kanamycin exposure to membrane defects, solute leakage, loss of turgor and cytoplasmic condensation ^29^. Although ampicillin and kanamycin initiate damage through distinct primary mechanisms and exhibit distinct patterns of envelope disruption, our single-cell analyses reveal that OM destabilization precedes downstream cellular failure in both cases. Thus, OM destabilization may represent a common transition point during antibiotic killing, although the mechanisms linking distinct antibiotic-induced damage processes to OM failure remain unresolved. Future studies will be required to define how diverse damage processes are coupled to the mechanical destabilization of the OM.

The Gram-negative OM has traditionally been viewed as a permeability barrier that contributes to the intrinsic antibiotic resistance and limits antibiotic discovery ^44^. However, recent studies have established that the OM also functions as a load-bearing structure that contributes to cellular mechanics, resistance to deformation and maintenance of cell shape ^30,45^. Our findings extend this mechanical role to antibiotic tolerance by demonstrating that OM stiffness directly influences survival under antibiotic killing. Indentation-depth-resolved AFM revealed a preferential increase in OM stiffness in multidrug-tolerant cells rather than a uniform increase in whole-envelope rigidity. Although intact-cell AFM measurements integrate contributions from multiple envelope components, the depth dependence of this phenotype supports localized remodeling of OM mechanics. Consistent with the view that OM mechanics emerges from the coordinated organization of LPS, OM proteins and OM–PG interactions ^30,33^, divalent-cation chelation reduced both OM stiffness and antibiotic tolerance, implicating that LPS-mediated interactions contribute to the elevated OM stiffness. Future studies will be required to determine how individual envelope components are coordinated to generate elevated OM stiffness.

The identification of the *amn*^T^ allele provided a genetic entry point for uncovering the mechanical basis of multidrug tolerance. Amn is a cytoplasmic AMP nucleosidase involved in adenylate metabolism, and altered Amn activity has been linked to adaptation to environmental stress ^46^, but not previously to antibiotic tolerance or envelope mechanics. Although the R473C substitution reduced canonical AMP nucleosidase activity, neither loss of *amn* nor catalytic impairment reproduced multidrug tolerance, indicating that altered Amn catalysis alone is insufficient to explain the mechanical remodeling of the OM. Defining how genetic perturbations are translated into altered envelope mechanics will be essential for understanding the molecular pathways that remodel OM organization and mechanics.

In summary, our study establishes bacterial envelope mechanics as a physical determinant of the transition from antibiotic-induced damage to lethal cellular failure and provides a mechanistic basis for exploring envelope mechanics as a complementary avenue for antimicrobial intervention.

## Materials and Methods

### Bacterial strains and culture

Two wild-type *Escherichia coli* genetic backgrounds were used: KLY ^18^ and BW25113. Experimental evolution was performed in the KLY background, and the effect of the *amn*^T^ allele was tested in both KLY and BW25113 backgrounds. The Δ*amn* and Δ*lpp* deletions were examined in the KLY background for functional analysis. Wild-type Amn and Amn variants were expressed in the KLY Δ*amn* background.

Unless otherwise specified, strains were grown overnight in lysogeny broth (LB; 10 g/l NaCl) medium at 37°C with shaking at 220 rpm, diluted 100-fold into fresh LB medium and cultured under the same conditions for the indicated duration before use.

### Appearance-time analysis by ScanLag

Overnight cultures were diluted 100-fold into fresh LB and grown at 37°C for 24 h. Cultures were serially diluted in PBS and plated on LB agar at approximately 150–200 colonies per plate. Plates were incubated at 37°C in a custom ScanLag automated scanner-array system ^47^. Plate images were acquired every 20 min for three days. Colony appearance times were extracted from the time-series images by automated image analysis and used to generate appearance-time distributions for each strain.

### Antibiotic survival assay

Overnight cultures were diluted 100-fold into fresh LB and grown at 37°C for 18 h before β-lactam and quinolone killing assays or for 24 h before aminoglycoside and polymyxin killing assays. For time-kill assays, cultures were then diluted 100-fold into fresh LB containing 120 µg/ml kanamycin, 50 µg/ml gentamicin, 120 µg/ml ampicillin, 2 µg/ml ertapenem, 8 µg/ml norfloxacin, 8 µg/ml ciprofloxacin, 10 µg/ml colistin or 10 µg/ml polymyxin B. At the indicated time points, 1-ml aliquots were sampled and washed four times with PBS to remove residual antibiotic. Survivors were serially diluted in PBS and plated on LB agar. Colony-forming units (CFUs) were determined after incubation for 2 days to allow all colonies to appear. Survival was calculated as the CFU count after antibiotic treatment relative to that immediately before antibiotic exposure.

For end-point survival assays, exposure durations were selected from the second phase of the biphasic time-kill curves, in which changes in survival predominantly reflected antibiotic tolerance. Unless otherwise specified, survival was measured after 5 h of kanamycin treatment, 5 h of ampicillin treatment, 8 h of norfloxacin treatment or 2 h of colistin treatment at the concentrations specified above. Samples were processed as described for the time-kill assays.

For experiments examining chemical and physical perturbations of antibiotic tolerance, 18-h or 24-h cultures prepared as described above were collected by centrifugation and resuspended in an equal volume of aqueous sorbitol solution containing 0.3, 0.6, 0.9 or 1.5 M sorbitol. For EDTA perturbation, to control for osmolarity, cells were resuspended in 1.5 M sorbitol containing 1, 5 or 10 mM EDTA, with or without 10 mM MgCl_2_ or CaCl_2_. Cell suspensions were incubated at 37°C for 1 h, collected by centrifugation and washed once with PBS before antibiotic challenge. Cells were then diluted 100-fold into fresh LB containing the indicated antibiotics. At the indicated time points, survivors were serially diluted in PBS and plated on LB agar. CFUs were determined after incubation for 2 days, and survival was calculated relative to the CFU count immediately before antibiotic treatment.

### Antibiotic susceptibility assays

For disk-diffusion assays, overnight cultures were diluted to approximately 10^6^ CFU/ml, and 100 µl of the bacterial suspension was spread evenly onto LB agar using sterile glass beads. Plates were allowed to dry for 15 min. Filter-paper disks (6 mm in diameter) were cut from filter paper (Whatman, #1) and sterilized. Each disk was loaded with 10 µl of a 1 mg/ml kanamycin or ampicillin solution, corresponding to 10 µg antibiotic per disk, and placed in the center of each plate. Plates were incubated at 37 °C for 18 h, and antibiotic susceptibility was assessed by measuring the radius of the zone of inhibition.

For liquid growth inhibition assays, overnight cultures were diluted into fresh LB medium to a final inoculum of approximately 10^5^ CFU/ml. Antibiotics were prepared as twofold serial dilutions in LB in 96-well plates, and bacterial suspensions were added to a final volume of 200 µl per well. Plates were incubated at 37°C for 24 h. OD_600_ was measured using a Tecan Infinite 200 microplate reader. Growth was quantified as the blank-subtracted net increase in OD_600_ (ΔOD_600_). The MIC was defined as the lowest antibiotic concentration at which no detectable bacterial growth was detected.

### Kanamycin-based experimental evolutions

Experimental evolution was performed in the KLY background using cyclic antibiotic selection. Two selection schemes were used: kanamycin-only evolution and a modified kanamycin-centered evolution incorporating ampicillin counterselection. Parallel overnight cultures (0.5 ml each) were diluted 100-fold into 50 ml fresh LB before antibiotic exposure.

For kanamycin-only evolution, cultures were exposed to 120 µg/ml kanamycin for 3 h at 37°C with shaking. Cells were collected by centrifugation at 6,000 g for 5 min, washed four times with PBS to remove residual kanamycin and resuspended in 1 ml fresh LB. Survivors were regrown for approximately 20 h at 37°C with shaking before the next selection cycle.

For the modified kanamycin-centered evolution, cultures were first exposed to 120 µg/ml kanamycin and 120 µg/ml ampicillin for 3 h at 37°C with shaking. Antibiotics were then removed by centrifugation at 6,000 g for 5 min followed by four washes with PBS. Cell pellets were then resuspended in 10 ml LB medium containing 120 µg/ml ampicillin and incubated for an additional 5 h as a counterselection step. Cells were subsequently collected by centrifugation at 6,000 g for 5 min, washed twice with PBS and resuspended in 1 ml fresh LB. Survivors were regrown for approximately 38 h at 37°C with shaking before the next cycle.

Batch cultures from each cycle and individual clones isolated from evolved populations were stored as glycerol stocks at −80°C for subsequent phenotypic and genetic analyses. *C3A5a*, *C3A5b* and *C3A5c* were isolated as individual clones from the batch cultures collected after the fifth selection cycle.

### Whole-genome sequencing

Strains were grown overnight in LB, and genomic DNA was extracted using the TIANamp Bacteria DNA Kit (TIANGEN). Whole-genome sequencing was performed by Beijing Sinobiocore Biological Technology Co., Ltd. on a DNBSEQ-T7 platform using 150-bp paired-end reads, with an average coverage of 150×. Reads were mapped and sequence variants were identified using Geneious Prime v2024.0.7. Candidate variants were validated by PCR amplification followed by Sanger sequencing.

### CRISPR-Cas9 genome editing

Genome editing of *amn* and *lpp* was performed by CRISPR–Cas9-mediated recombineering as described previously ^48^. Briefly, the Cas9- and λ-Red-expressing plasmid (p15A-pBAD-Cas9-pT5-Redγβα) and the sgRNA plasmid (pSC101-pBAD-sgRNA-Donor) carrying the donor sequence were co-transferred into wild-type or *amn*^T^ cells. Transformants were selected on LB agar containing 100 µg/ml kanamycin, 100 µg/ml ampicillin and 10 mg/ml glucose and recovered in fresh LB containing the same antibiotics and glucose at 30°C for 2 h. Protein expression was sequentially induced with 1 mM IPTG for 1 h and 20 mM L-arabinose for 3 h. Cultures were then diluted 1,000-fold into fresh LB and plated on LB agar containing 100 µg/ml kanamycin, 100 µg/ml ampicillin and 20 mM L-arabinose. Genome editing was verified by PCR and Sanger sequencing. Editing plasmids were subsequently cured by growth in LB at 40°C for 12 h.

### Protein purification

Overnight cultures of *E. coli* BL21(DE3) expressing N-terminal 3×FLAG-tagged wild-type or mutant Amn proteins were diluted 100-fold into fresh LB containing 100 µg/ml kanamycin and grown at 37°C with shaking to an OD_600_ of ∼0.6–0.8. Protein expression was induced with 0.5 mM IPTG, followed by incubation at 16°C for 20 h. Cells were collected by centrifugation (4,000 g, 20 min), resuspended in lysis buffer (50 mM Tris-HCl pH 7.4, 150 mM NaCl, 1 mM EDTA, 0.1% Triton X-100, 100 µg/ml lysozyme), and disrupted by sonication on ice (2 s on, 2 s off, 20 min total). After centrifugation (12,000 g, 20 min, 4°C), the supernatant was loaded onto Anti-FLAG Affinity Gel (Yeasen), and the resin was washed three times with wash buffer (50 mM Tris-HCl pH 7.4 containing 150 mM NaCl). Bound proteins were eluted with 100 µg/ml 3×FLAG peptide in wash buffer. Purified proteins were analyzed by SDS– PAGE and Coomassie staining, quantified by BCA assay and stored at −80°C until use.

### Xanthine oxidase-coupled Amn activity assay

The in vitro activity of Amn variants was measured using a xanthine oxidase-coupled assay modified from previously described methods ^49,50^. In brief, adenine released from AMP by Amn was oxidized by xanthine oxidase, coupled to the reduction of iodonitrotetrazolium chloride (INT) to formazan, which absorbs at 500 nm. Reactions were performed in a final volume of 100 µl containing 1 mM INT, 0.2 units of xanthine oxidase (MCE), 400 µM Mg–ATP, 10 µl DMSO and 0–1 mM AMP in reaction buffer (104 mM KCl, 50 mM Tris-HCl pH 8.0). Reaction mixtures without Amn were used to determine background absorbance. Reactions were initiated by addition of purified Amn to a final concentration of 100 nM and incubated at 37°C for 20 min. Absorbance at 500 nm was measured using a Tecan Infinite 200 microplate reader.

Adenine production was quantified using an adenine standard curve generated under the same detection conditions. Adenine standards (0–1 mM) was subjected to the xanthine oxidase-coupled reaction in the absence of AMP and Amn, and the resulting adenine–A500 curve was used to convert the absorbance values to adenine concentrations. Amn activity was calculated from the amount of adenine generated over 20 min and expressed as μM/min.

### ROS-associated fluorescence assay

Overnight cultures were diluted 100-fold in fresh LB and grown at 37°C for 24 h. Cultures were then diluted 100-fold in fresh LB, incubated with 10 µM dihydroethidium (DHE; Reactive Oxygen Species Assay Kit, Applygen) for 30 min before treatment with 120 µg/ml kanamycin, 120 µg/ml ampicillin, 8 µg/ml norfloxacin, 10 µg/ml colistin or 10 mM H_2_O_2_ for 2 h at 37 °C. Fluorescence was measured using a Tecan Infinite 200 microplate reader with excitation and emission wavelengths of 535 nm and 610 nm, respectively. Wells without bacteria were used to determine background fluorescence and OD_600_. After background subtraction, ROS-associated fluorescence was normalized to cell density as: normalized fluorescence = (fluorescence of treated sample – fluorescence of blank) / (OD_600_ of treated sample – OD_600_ of blank)

### Protein synthesis detection

Protein synthesis was monitored from the accumulation of chromosomal expressing cytoplasmic YFP fluorescence. Overnight cultures were diluted 100-fold in fresh LB medium and grown at 37°C for 24 h. Cultures were then diluted 100-fold into fresh LB medium, treated with 120 µg/ml kanamycin incubated at 37 °C. Cytoplasmic YFP fluorescence was measured every 10 min for 24 h using a Tecan Infinite 200 microplate reader with excitation and emission wavelengths of 510 nm and 527 nm, respectively.

### SOS activation detection

SOS activation was measured using a pET28-derived tetracycline-resistant reporter plasmid carrying mCherry under the control of the *sulA* promoter. KLY wild-type and *amn*^T^ cells were transformed with P*sulA*-mCherry reporter plasmid and cultured into fresh LB containing 10 µg/ml tetracycline. Overnight cultures were diluted 100-fold into fresh LB and grown at 37°C for 24 h. Cultures were then diluted 100-fold into fresh LB and treated with norfloxacin at 37°C for 8 h. mCherry fluorescence (excitation, 587 nm; emission, 610 nm) and OD_600_ were measured using a microplate reader (Tecan Spark). Wells without bacteria were used to determine background fluorescence and OD_600_. Fluorescence was normalized to cell density after background subtraction using the equation described above.

### NPN/PI membrane-permeability assay

Outer- and inner-membrane permeability were assessed using N-phenyl-1-naphthylamine (NPN) and propidium iodide (PI), respectively. Overnight cultures were diluted 100-fold into fresh LB and grown at 37°C for 18 h for ampicillin assays and 24 h for kanamycin assays. Cultures were then diluted 100-fold into fresh LB supplemented with or without 120 µg/ml ampicillin or 120 µg/ml kanamycin. At the indicated time points, cells were collected, washed once by PBS and resuspended in 100 µl of 10 mM HEPES buffer (pH 7.0) containing 10 µM NPN and 10 µM PI. After incubation for 20 min, NPN fluorescence was measured at an excitation wavelength of 350 nm and an emission wavelength of 420 nm, and PI fluorescence was measured at an excitation wavelength of 535 nm and an emission wavelength of 617 nm using a microplate reader (Tecan Spark). HEPES buffer containing NPN and PI but no bacteria was processed in parallel as a background control. Fluorescence signals were normalized to OD_600_ after background subtraction using the equation described above.

### Time-lapse fluorescence microscopy for antibiotic killing

To simultaneously monitor periplasmic and cytoplasmic responses during antibiotic treatment, KLY wild-type and *amn*^T^ cells constitutively expressing chromosomal YFP were transformed with a pET28-derived, tetracycline-resistant plasmid encoding mCherry fused to the DsbA signal peptide (_ss_DsbA– mCherry) for periplasmic localization. The plasmid was maintained with tetracycline, and mCherry expression was induced by IPTG.

Overnight cultures were diluted 100-fold into fresh LB containing 10 µg/ml tetracycline and 500 µM IPTG and grown at 37°C for 18 h for ampicillin imaging or 24 h for kanamycin imaging. For imaging of EDTA treated *amn*^T^ cells, cultures were collected by centrifugation and resuspended in an equal volume of 1.5 M sorbitol containing 10 mM EDTA for 1 h at 37°C. Cells were then collected by centrifugation, washed once with PBS and resuspended in fresh LB before imaging. For imaging, 1 µl of bacterial culture was spotted onto the center of a confocal dish (Cellvis). A 1 × 1 cm agarose pad prepared from 0.75% LB agarose containing 120 µg/ml ampicillin or 120 µg/ml kanamycin was gently placed over the cells, and excess liquid around the pad was removed before imaging.

Time-lapse imaging was performed at 37°C using an LSM980 (Carl Zeiss, Jena) confocal microscope equipped with Airyscan 2 module and a Plan-Apochromat 100x/1.4 Oil DIC M27 (Zeiss). YFP and mCherry were excited at 514nm and 594nm, respectively, and fluorescence was collected over over 410nm-579nm and 410nm-694nm. Image acquisition was controlled using ZEN Blue v.3.9. Images were acquired with 1.2× zoom, 0.075um per pixel, 8 bits per pixel and without averaging, result in an imaging size 70.71 µm × 70.71 µm, 947 pixels × 947 pixels. Images were acquired every 5 min for 3 h for ampicillin and 5 h for kanamycin. Raw images were processed using the 2D AiryScan processing function in ZEN Blue and exported for further analysis in Fiji v.2.16.0/1.54p. Cell morphology, focal redistribution of periplasmic mCherry, cytoplasmic bulging or condensation, lysis and severe deformation were quantified from individual-cell time-lapse sequences.

### AFM measurements of cell stiffness

AFM samples were prepared as previously described ^30^, with minor modifications. Briefly, 8-mm circular glass coverslips were boiled in 2% Micro-90 detergent, rinsed with ethanol and dried by heating. On the day of measurement, coverslips were attached to 15-mm iron discs using 3M Super Glue, coated with 10 µl of 0.01% poly-L-lysine (Sigma), incubated for 90 min and then air-dried before use.

Overnight *E. coli* cultures were diluted 100-fold into fresh LB and grown at 37°C for 24 h. For untreated samples, stationary-phase cultures were diluted 100-fold into fresh LB and grown at 37°C for 1.5–2 h, corresponding to an increase in OD_600_ (ΔOD_600_) of approximately 0.02, measured using a Tecan Infinite 200 microplate reader. For chemical and physical perturbation experiments, treated cells were collected by centrifugation, washed once with PBS, diluted into fresh LB and grown under the same conditions for 1.5–2 h to a ΔOD_600_ of approximately 0.02. Cells were collected by centrifugation (6,000 g, 2 min) and resuspended in PBS to approximately 10^8^ CFU/ml. A 100 µl aliquot of cell suspension was applied to poly-L-lysine-coated coverslips and incubated at room temperature for 15 min to allow cell attachment. Coverslips were rinsed three times with PBS to remove unattached cells and covered with 50 µl PBS during AFM measurement.

AFM experiments were performed on a Bruker Multimode 8-HR (Bruker) with PFQNM-LC-V2 (Bruker) cantilevers. The nominal spring constant of each cantilever was provided by the manufacturer. Before measurements, the deflection sensitivity was calibrated using the thermal noise method. For each cell, force curves were recorded in a scan size of 500 × 500 nm at a resolution of 64 × 64 pixels at 1 Hz scan rate. Force curves were analyzed using NanoScope Analysis v.3.0 (Bruker). Baseline correction of the extend curve was performed using 10–80% of the curve. For mechanical analysis, extend curves were fitted using the Hertz contact model ^51^ over the 20–60% maximum force range, assuming a spherical tip with a radius of 70 nm and a Poisson’s ratio of 0.3. For each cell, the mean Young’s modulus (*E*_AFM_), used as a measure of stiffness, was calculated across all force curves within the recorded region. Indentation depth was defined as the displacement from the contact point on the extend curve to the point of maximum displacement at the maximum load, and the mean indentation depth for each cell was calculated across all force curves within the recorded region. In all cases, individual samples were measured for no more than 90 min.

### Statistical and quantitative stiffness–survival analysis

Statistical analyses were performed using GraphPad Prism v.10.5.0. Unless otherwise specified, data are presented as mean ± s.e.m. Biological replicate numbers and statistical tests are indicated in the corresponding figure legends. Comparisons between two groups were performed using two-sided unpaired Student’s t-tests. Comparisons involving multiple groups or conditions were performed using one-way or two-way ANOVA followed by Bonferroni correction for multiple comparisons. No statistical method was used to predetermine sample size. Exact n values are provided in the figure legends. *P* < 0.05 was considered statistically significant.

To quantify the relationship between cell-envelope stiffness and antibiotic survival, survival rates were log_10_-transformed before analysis. For each experimental condition, Young’s modulus (*E*_AFM_) was paired with the corresponding mean survival measured under the same condition. The stiffness–survival relationship was analyzed independently for kanamycin, ampicillin, norfloxacin and colistin using ordinary least-squares linear regression. Regression models were fitted as

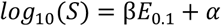

where *S* is the log_10_-transformed survival rate, and *E*_0.1_ represents stiffness expressed in units of 0.1 MPa. The regression coefficient *β* represents the change in log_10_ survival associated with a 0.1-MPa increase in stiffness, and *α* is the intercept. For each antibiotic, *β* estimates, 95% confidence intervals, R^2^ values and *P* values were obtained from the fitted regression model.

For comparisons between perturbation types, chemical and physical perturbation datasets were analyzed independently using the same regression procedure. Stiffness coefficients derived from each dataset were compared on the basis of their *β* estimates and 95% confidence intervals. All regression analyses were performed in GraphPad Prism v.10.5.0.

## Acknowledgement

We thank Lin Ge and Yan Zhang of the Center of Pharmaceutical Technology at Tsinghua University, for their assistance with Multimode 8-HR (Bruker) AFM and LSM980 (Carl Zeiss, Jena) confocal microscope. The CRISPR/Cas9-based genome editing tool were generously donated by Huo’s lab, School of Life Science, Beijing Institute of Technology. We appreciate Nathalie Balaban’s lab at Racah Institute of Physics, The Hebrew University of Jerusalem, for providing the *E. coli* KLY strain. We thank Y. Li (School of Life Sciences, Southern University of Science and Technology) and K. Zhu (College of Veterinary Medicine, China Agricultural University) for helpful discussions and valuable suggestions.

## Author Contributions

N.X., and J.-F.L. conceived the project, designed the study and wrote the paper. N.X. designed and performed the evolution experiments, tolerance detections, and AFM measurements. B.Z. performed genome editing experiments and conducted bioinformatic analyzes. N.X. interpreted the results. J.-F.L. supervised the project.

## Data and Materials Availability

All data needed to evaluate the conclusions in the paper are present in the paper and/or the Supplementary Materials.

