## Supplementary File for "Outer membrane stiffness gates multidrug tolerance"

**The PDF file includes:**

Extended Data Fig. 1 to 9

Extended Data Table 1 to 2

**Other Supplementary Material for this manuscript includes the following:**

Supplementary Movie 1 to 6

### Extended Data Figures

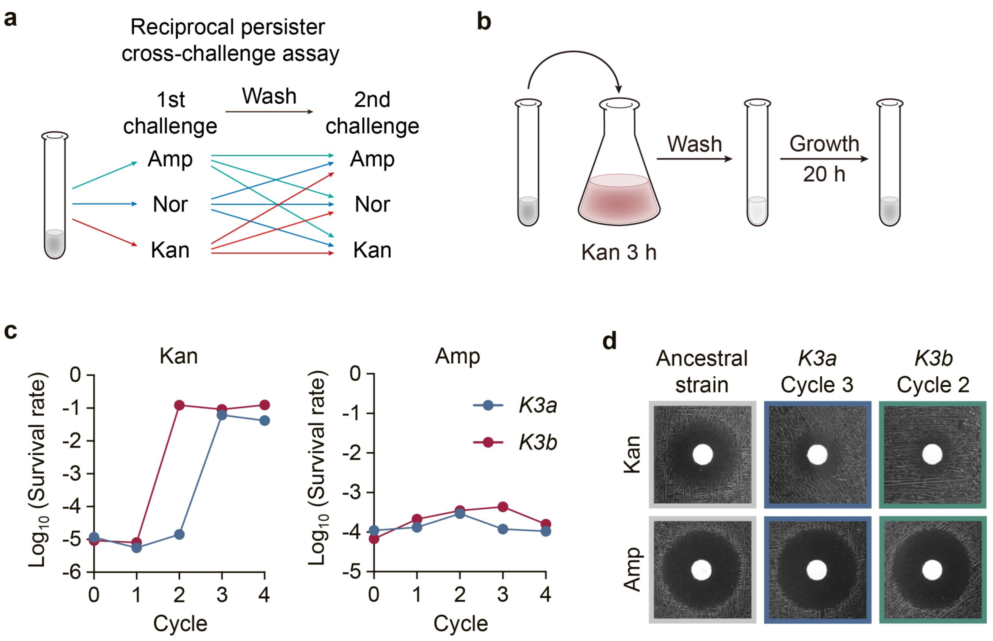


**Extended Data Fig. 1 | Persister cross-challenge assay and kanamycin-only evolution. a**, Schematic of the reciprocal persister cross-challenge assay. Cells surviving primary antibiotic treatment were isolated, washed and re-challenged with the indicated antibiotics. Kanamycin, Kan; ampicillin, Amp; norfloxacin, Nor. **b**, Schematic of kanamycin-only evolution. Overnight cultures were diluted 100-fold in fresh LB medium containing 120 μg/ml kanamycin and treated for 3 h. After kanamycin washout, survivors were resuspended in fresh LB medium and allowed to regrow for 20 h. **c**, Survival of independently evolved *K3a* and *K3b* batch cultures at each cycle after 5 h treatment with kanamycin or ampicillin (each at 120 μg/ml). **d**, Disk diffusion assays of the ancestral strain and representative clones isolated from *K3a* and *K3b* evolved populations using 10 μg kanamycin or ampicillin disks.

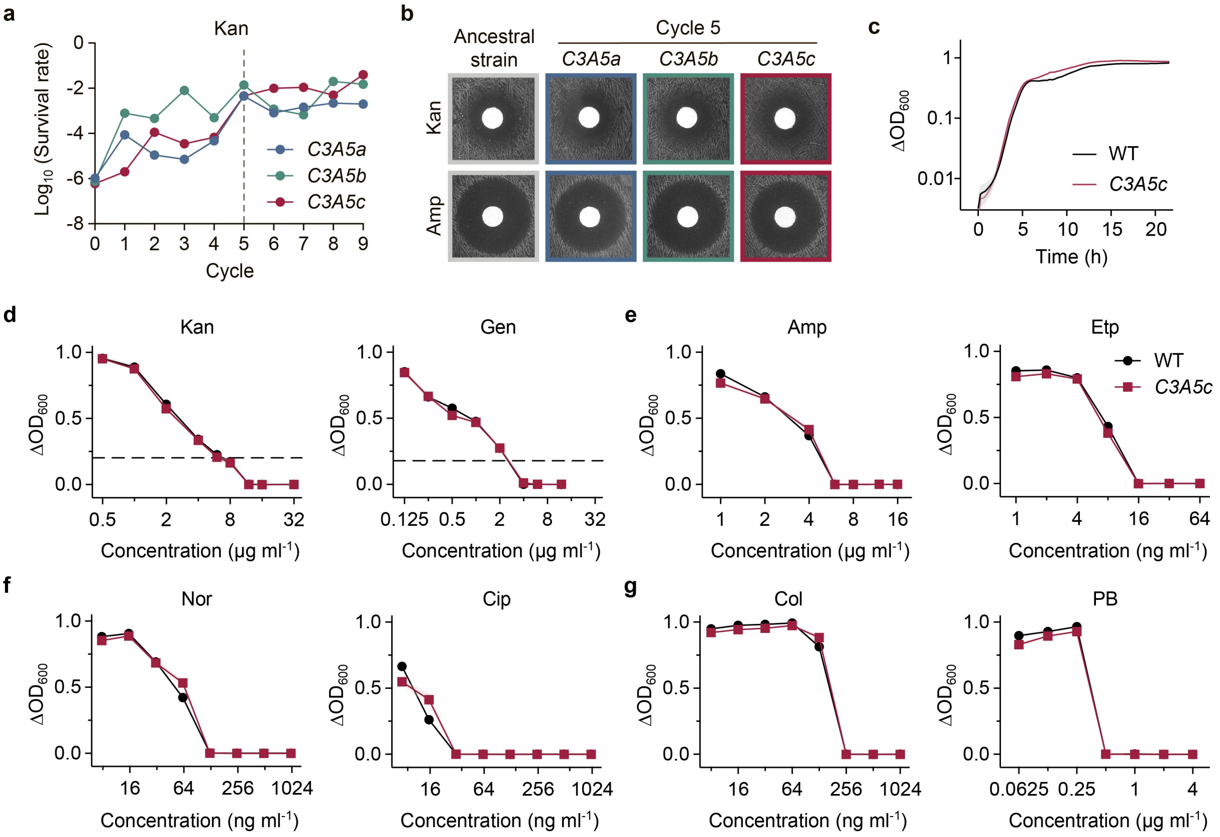


**Extended Data Fig. 2 | Kanamycin-centered evolution and characterization of evolved tolerant clones.** **a**, Survival of independently evolved batch cultures *C3A5a*, *C3A5b* and *C3A5c* at each cycle after 5 h treatment with 120 μg/ml kanamycin. **b**, Disk diffusion assays of the ancestral strain and evolved clones isolated from *C3A5a–c* populations using 10 μg kanamycin or ampicillin disks. **c**, Growth curves of wild-type (WT) and *C3A5c* cells in antibiotic-free fresh LB medium. **d**–**g**, Minimum inhibitory concentration (MIC) measurements of wild-type and *C3A5c* cells against aminoglycosides kanamycin and gentamicin (Gen; **d**), β-lactams ampicillin and ertapenem (Etp; **e**), quinolones norfloxacin and ciprofloxacin (Cip; **f**), or polymyxins colistin (Col) and polymyxin B (PB; **g**). Growth inhibition was monitored by $OD_{600}$ measurements after antibiotic exposure for 24 h. The dashed line in **e** indicates the $\Delta OD_{600}$ threshold used to determine aminoglycoside MICs. Data in **c**–**g** are mean ± s.e.m. from three biological replicates.

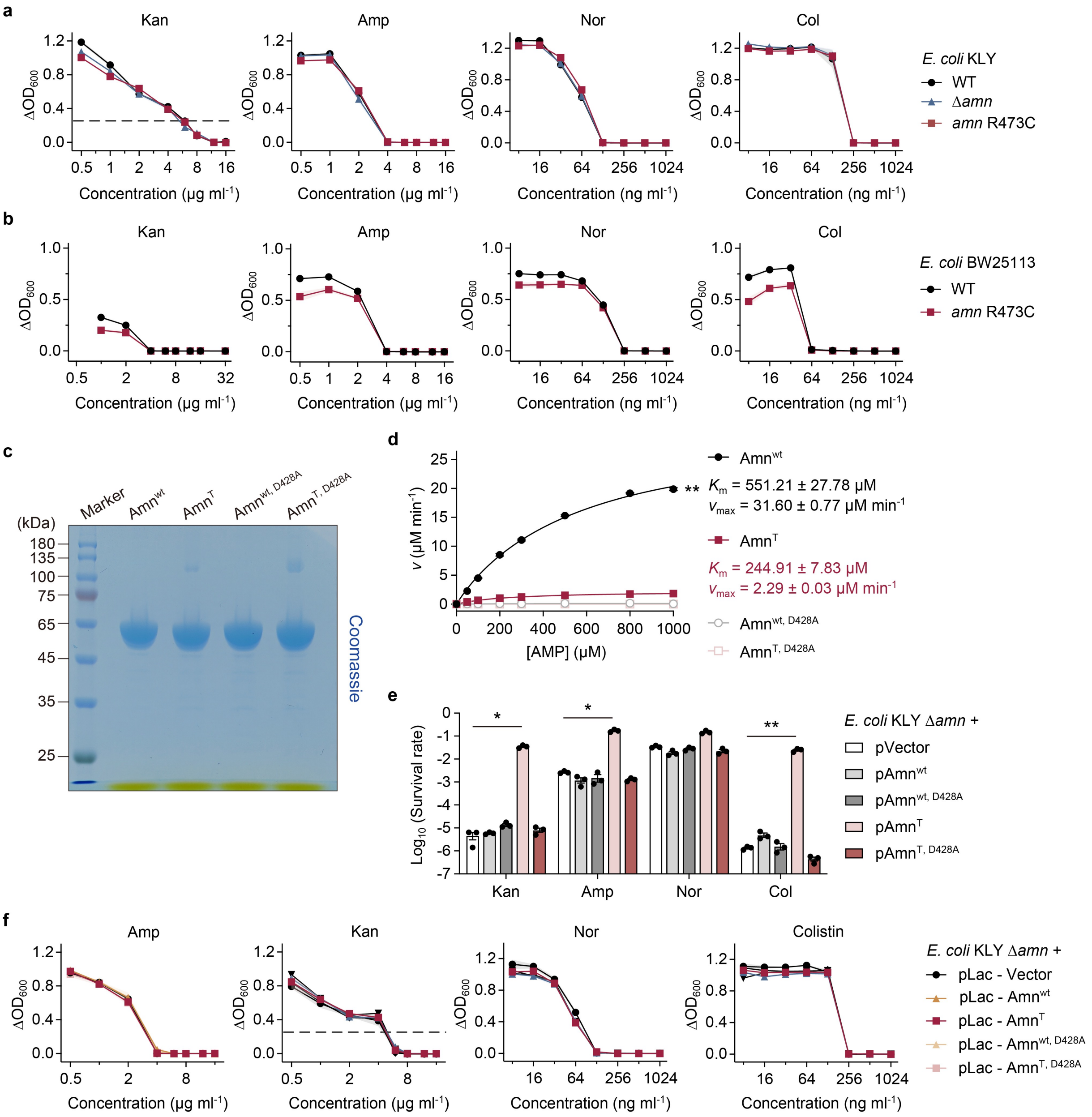


**Extended Data Fig. 3 | *amn*$^{T}$-mediated multidrug tolerance does not result from loss of Amn catalytic activity.** **a**, **b**, MIC measurements of wild-type, *amn*-deletion (Δ*amn*) and reconstructed *amn* R473C (*amn*$^{T}$) strains in *E. coli* KLY (**a**) and BW25113 (**b**) backgrounds against the indicated antibiotics. **c**, SDS–PAGE analysis of purified FLAG-tagged Amn, Amn R473C (Amn$^{T}$) and Amn D428A proteins used for enzymatic assays. **d**, Xanthine oxidase-coupled assay of AMP nucleosidase activity of purified Amn, Amn R473C and Amn D428A proteins. Adenine production was quantified using an adenine standard curve generated under the same reaction conditions. **e**, Survival of *E. coli KLY* Δ*amn* strains expressing empty vector, wild-type Amn, Amn R473C or Amn D428A following kanamycin (120 μg/ml, 5 h), ampicillin (120 μg/ml, 5 h), norfloxacin (8 μg/ml, 8 h) or colistin (10 μg/ml, 2 h) treatment. **f**, MIC measurements of *E. coli* KLY Δ*amn* strains expressing wild-type Amn, Amn R473C or Amn D428A. Data in **a**, **b** and **d**–**f** are mean ± s.e.m. from three biological replicates. Statistical significance was determined using a two-sided unpaired Student's t-test (**d**) or two-way ANOVA with Bonferroni correction (**e**); ns, not significant; *$P < 0.05$; **$P < 0.01$.

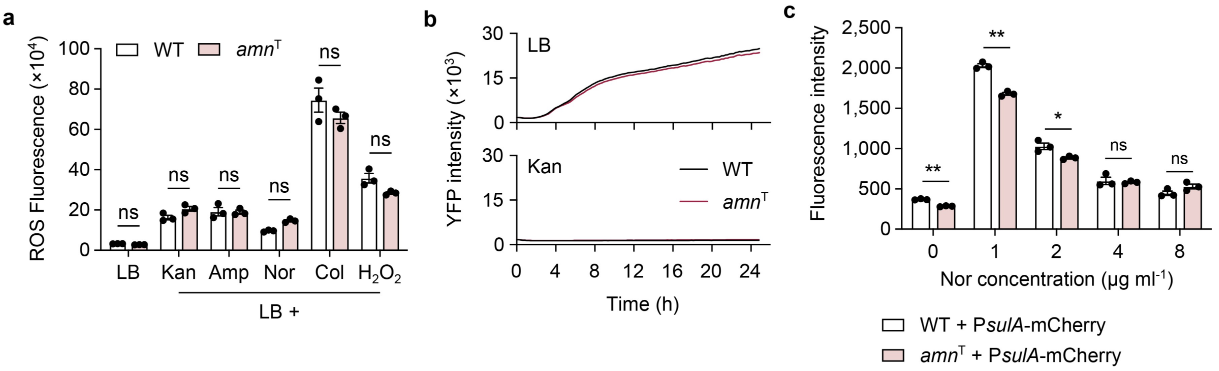


**Extended Data Fig. 4 | *amn*ᵀ does not broadly suppress canonical intracellular responses to antibiotic treatment.** **a**, ROS-associated fluorescence of wild-type and *amn*ᵀ cells in LB medium or following treatment with kanamycin (120 μg/ml), ampicillin (120 μg/ml), norfloxacin (8 μg/ml), colistin (10 μg/ml) or $H_2O_2$ (5 mM) for 3 h. ROS-associated fluorescence was measured using 10 μM dihydroethidium and normalized to cell density after background subtraction. **b**, Cytoplasmic YFP fluorescence of wild-type and *amn*ᵀ cells during growth in LB medium or following treatment with kanamycin (120 μg/ml). **c**, SOS reporter activity in wild-type and *amn*ᵀ cells carrying the P*sulA*-mCherry reporter following treatment with the indicated concentrations of norfloxacin. Fluorescence was normalized to cell density after background subtraction. Data are mean ± s.e.m. from three biological replicates. Statistical significance was determined using two-way ANOVA with Bonferroni correction; ns, not significant; $^{*}P < 0.05$; $^{**}P < 0.01$.

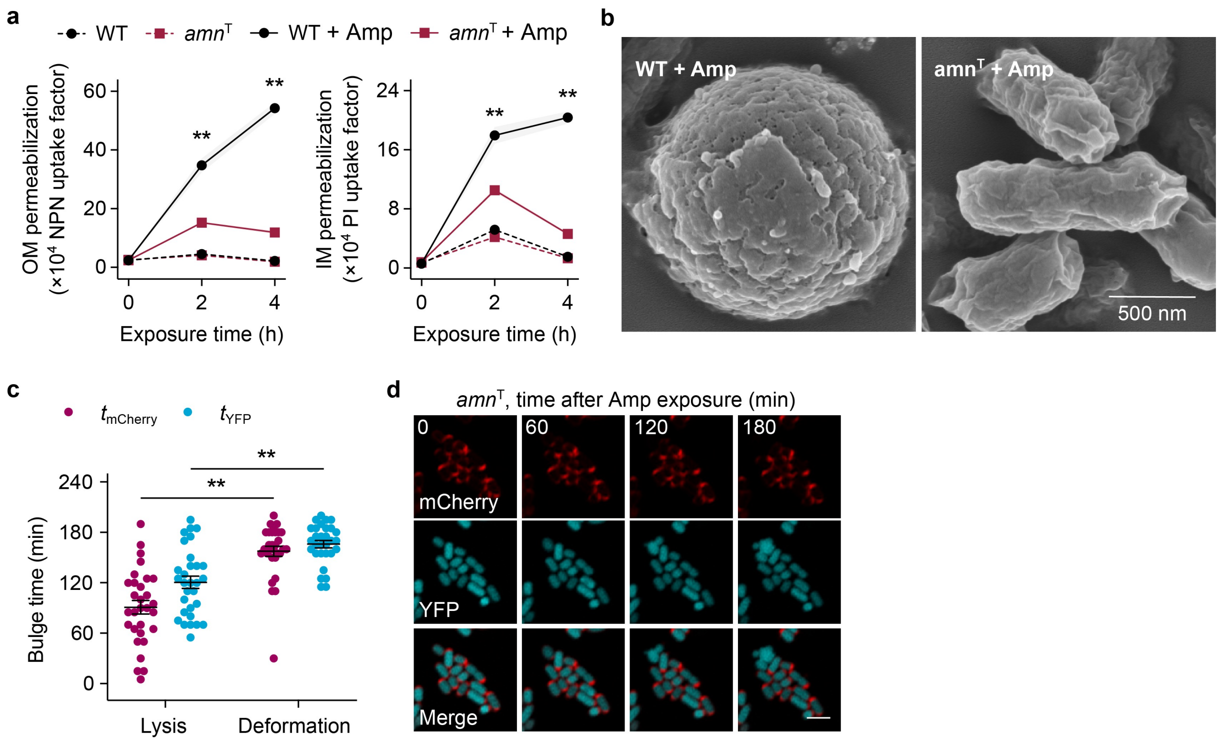


**Extended Data Fig. 5 | *amn*$^{T}$ attenuates ampicillin-induced membrane permeabilization and envelope deformation. a**, Outer-membrane (OM) and inner-membrane (IM) permeabilization of wild-type and *amn*$^{T}$ cells during ampicillin exposure, measured by N-phenyl-1-naphthylamine (NPN) and propidium iodide (PI) uptake, respectively ($n$ = 3 biological replicates per group). **b**, Representative scanning electron microscopy images of wild-type and *amn*$^{T}$ cells following ampicillin treatment. Scale bar, 500 nm. **c**, Onset times of periplasmic mCherry- and cytoplasmic YFP-associated morphological events in wild-type cells undergoing lysis or non-lytic deformation during ampicillin exposure ($n = 20$ cells per group). **d**, Representative time-lapse fluorescence images of *amn*$^{T}$ cells during ampicillin exposure, showing periplasmic $_{ss}$DsbA–mCherry (red) and cytoplasmic YFP (cyan). Scale bar, 2 µm. Ampicillin was used at 120 µg/ml. Data in **a** and **c** are mean ± s.e.m. Statistical significance was determined using two-way ANOVA with Bonferroni correction; $^{**}P < 0.01$.

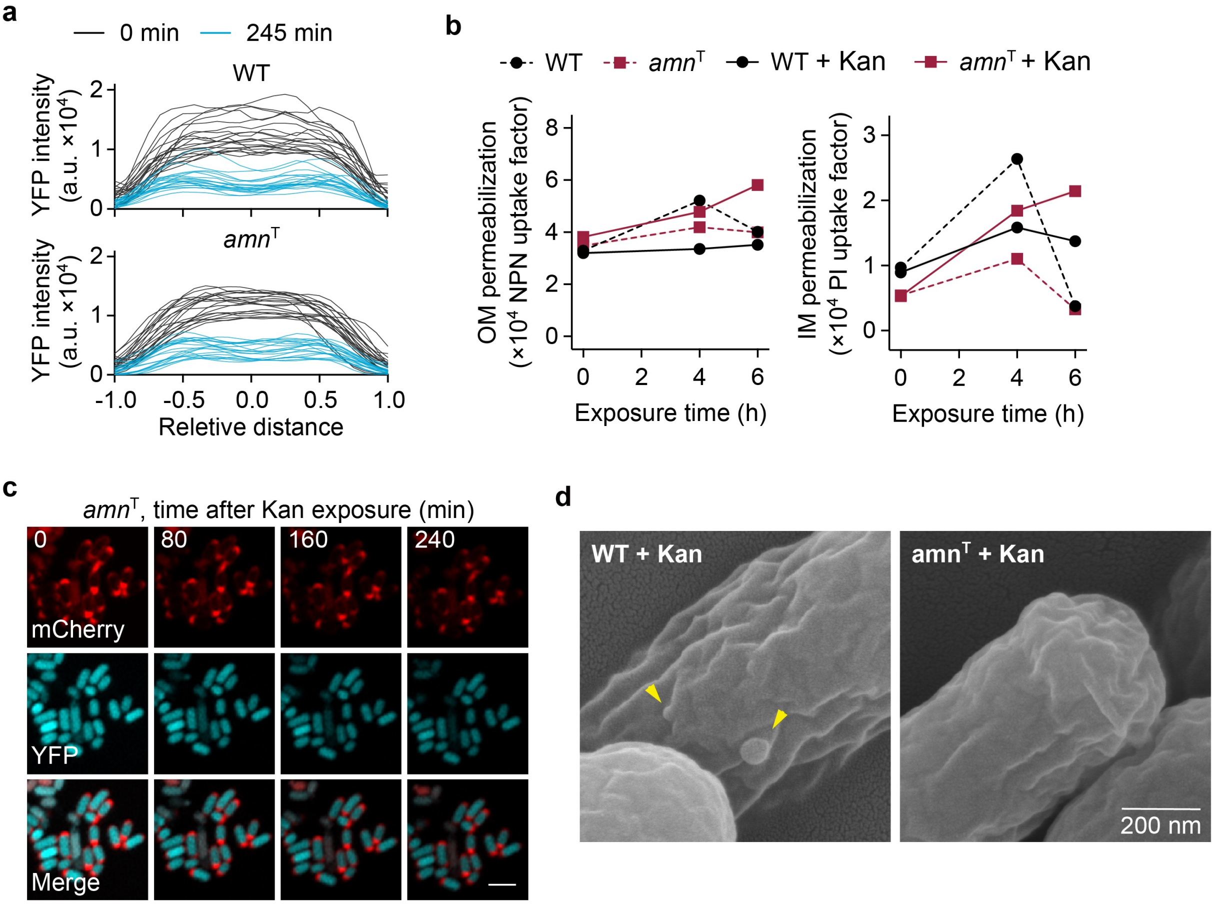


**Extended Data Fig. 6 | *amn*$^T$ suppresses kanamycin-induced envelope destabilization without preventing cytoplasmic condensation.** **a**, Longitudinal cytoplasmic YFP fluorescence-intensity profiles of individual wild-type and *amn*$^T$ cells before and after 245 min of kanamycin exposure (*n* = 20 cells per group). Cell length was normalized from −1 to 1. **b**, Outer-membrane and inner-membrane permeabilization of wild-type and *amn*$^T$ cells during kanamycin exposure, measured by NPN and PI uptake, respectively (*n* = 3 biological replicates per group). Data are mean ± s.e.m. **c**, Representative time-lapse fluorescence images of *amn*$^T$ cells during kanamycin exposure, showing periplasmic $_{ss}$DsbA–mCherry (red) and cytoplasmic YFP (cyan). Scale bar, 2 μm. **d**, Representative scanning electron microscopy images of wild-type and *amn*$^T$ cells following kanamycin treatment. Arrowheads indicate focal OM protrusions in wild-type cells. Data in **b** are mean ± s.e.m. Scale bar, 200 nm. Kanamycin was used at 120 μg/ml.

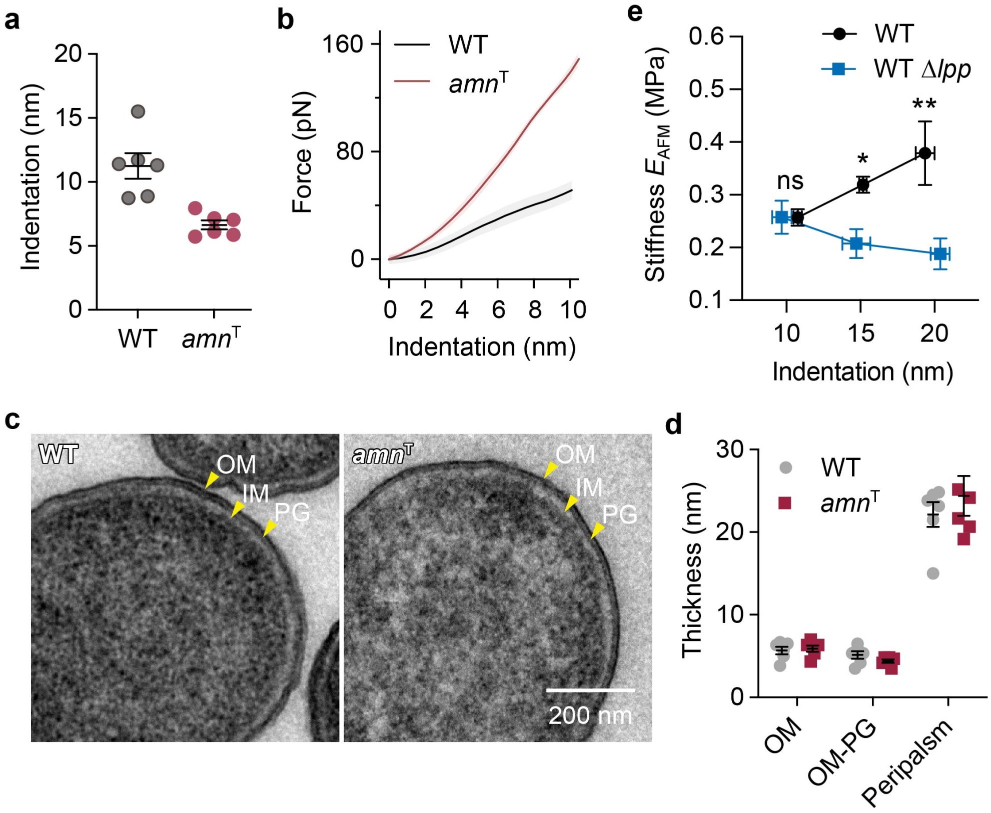


**Extended Data Fig. 7 | Shallow AFM indentation preferentially resolves OM-associated mechanics.** **a**, Indentation depths of wild-type and *amn*$^{T}$ cells under the force conditions used for shallow atomic force microscopy (AFM) measurements ($n$ = 6 cells per group). **b**, Force–indentation curves of wild-type and *amn*$^{T}$ cells within the initial 10-nm indentation range ($n$ = 10 cells per group). **c**, Representative transmission electron microscopy (TEM) images of wild-type and *amn*$^{T}$ cells showing the organization of the outer membrane, peptidoglycan (PG) and inner membrane. Scale bar, 200 nm. **d**, Thickness measurements of envelope structures in wild-type and *amn*$^{T}$ cells ($n$ = 6 cells per group). **e**, Stiffness $E_{AFM}$ of wild-type and *lpp*-deleted (Δ*lpp*) cells measured at the indicated indentation depths ($n$ = 6 cells per group). Data are mean ± s.e.m. Statistical analysis was performed using two-way ANOVA with Bonferroni correction; ns, not significant; *$P < 0.05$; **$P < 0.01$.

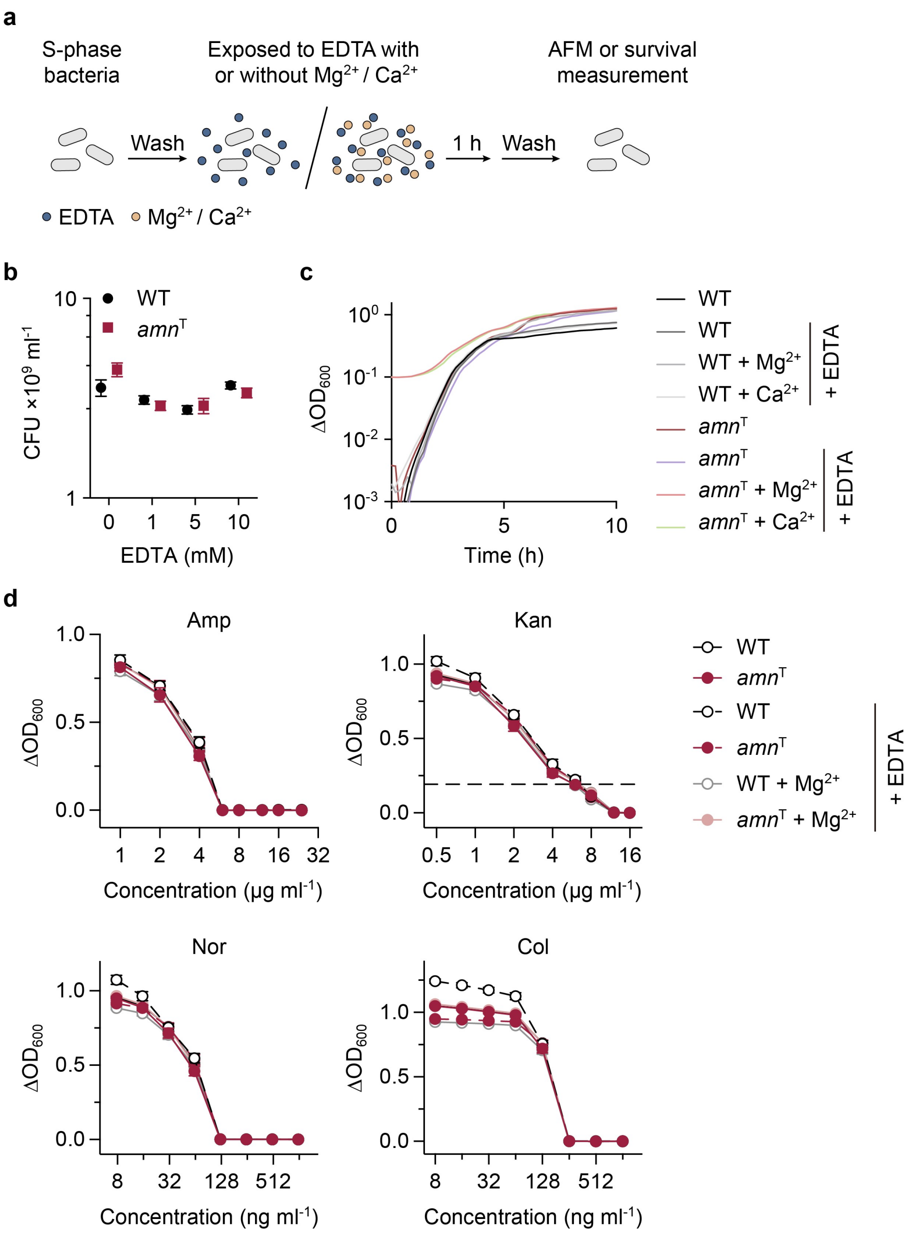


**Extended Data Fig. 8 | Transient EDTA perturbation does not alter cell viability, growth or antibiotic susceptibility. a**, Schematic of the transient EDTA perturbation. Stationary-phase cells were washed and exposed to EDTA for 1 h, with or without 10 mM $MgCl_2$ or $CaCl_2$ supplementation, followed by washout before AFM or antibiotic survival measurements. **b**, Colony-forming units (CFUs) of wild-type and *amn*$^T$ cells after treatment with the indicated concentrations of EDTA for 1 h. **c**, Growth curves of wild-type and *amn*$^T$ cells following 10 mM EDTA treatment, with or without 10 mM $MgCl_2$ or $CaCl_2$ supplementation. Growth was monitored by changes in $OD_{600}$. **d**, MIC measurements of wild-type and *amn*$^T$ cells following 10 mM EDTA treatment, with or without 10 mM $MgCl_2$ or $CaCl_2$ supplementation, against ampicillin, kanamycin, norfloxacin or colistin. Growth inhibition was monitored by $OD_{600}$ measurements after antibiotic exposure for 24 h. The dashed line indicates the $\Delta OD_{600}$ threshold used to determine the kanamycin MIC. Data are mean ± s.e.m. from three biological replicates.

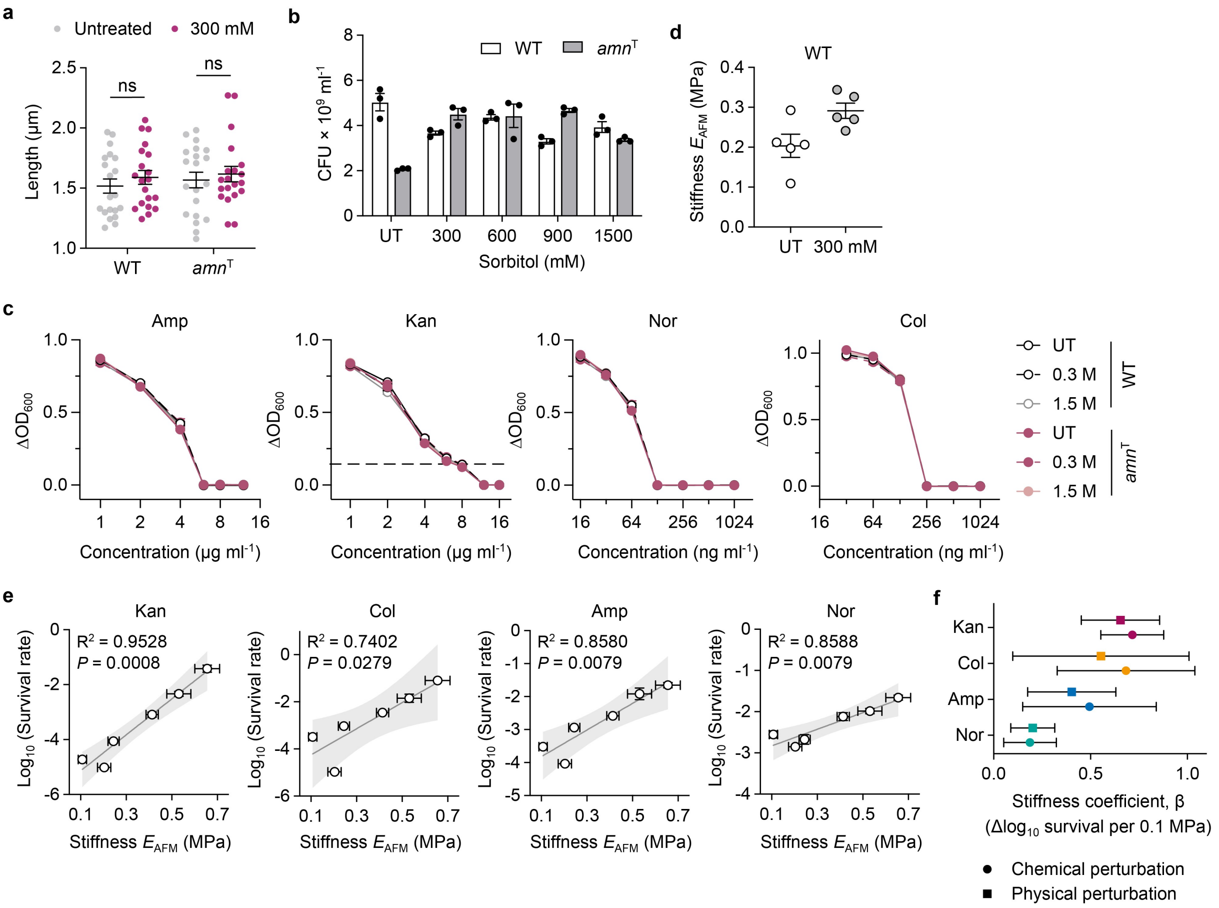


**Extended Data Fig. 9 | Independent osmotic perturbation supports the quantitative stiffness–survival relationship. a**, Cell length of wild-type and *amn*$^{T}$ cells before (UT) and after treatment with 300 mM sorbitol for 1 h (*n* = 20 cells per group). **b**, CFUs of wild-type and *amn*$^{T}$ cells before and after treatment with the indicated sorbitol concentrations for 1 h (*n* = 3 biological replicates per group). **c**, MIC measurements of wild-type and *amn*$^{T}$ cells following treatment with the indicated sorbitol concentrations, against ampicillin, kanamycin, norfloxacin or colistin (*n* = 3 biological replicates per group). Growth inhibition was monitored by $OD_{600}$ measurements after antibiotic exposure for 24 h. The dashed line indicates the $\Delta OD_{600}$ threshold used to determine the kanamycin MIC. **d**, Stiffness of wild-type cells before and after treatment with 300 mM sorbitol for 1 h (*n* = 5 cells per group). **e**, Relationship between stiffness and antibiotic survival across genetic and physical perturbations for kanamycin, colistin, ampicillin and norfloxacin. Solid lines indicate linear regression fits, with $R^2$ and *P* values shown. **f**, Comparison of stiffness coefficients (β) derived independently from chemical and physical perturbations for each antibiotic. Coefficients represent the change in $\log_{10}$ survival associated with a 0.1-MPa increase in stiffness. Error bars indicate 95% confidence intervals. Data are mean ± s.e.m. Statistical analysis in **a** was performed using two-way ANOVA with Bonferroni correction; relationships in **e** and **g** were assessed by linear regression; ns, not significant.

**Extended Data Table 1 | MICs and bactericidal treatment concentrations of antibiotics used for *E. coli* KLY.**

| Class | Antibiotic | MIC (μg/ml) | Killing concentration (μg/ml) | Killing concentration (× MIC) |
|---|---|---|---|---|
| β-lactam | Ampicillin | 6 | 120 | 20 |
| | Ertapenem | 0.016 | 2 | 125 |
| Aminoglycoside | Kanamycin | 6 | 120 | 20 |
| | Gentamicin | 2 | 50 | 25 |
| Quinolone | Norfloxacin | 0.125 | 8 | 64 |
| | Ciprofloxacin | 0.03 | 8 | 267 |
| Polymyxin | Colistin | 0.256 | 10 | 39 |
| | Polymyxin B | 0.5 | 10 | 20 |

**Extended Data Table 2 | Mutations detected by whole-genome sequencing and verified by Sanger sequencing.**

| Strain | Genomic position | Mutation | Amino acid substitution | Gene | Annotation |
|---|---|---|---|---|---|
| *C3A5a* | 2996764 -2996832 | $\Delta$69 bp | NA | NA | |
| | 3038716 | T>C | S102P (TCC→CCC) | *trbA* | Conjugal transfer protein TrbA |
| *C3A5b* | 3038716 | T>C | S102P (TCC→CCC) | *trbA* | Conjugal transfer protein TrbA |
| *C3A5c* ($amn^{\mathrm{T}}$) | 2032182 | C>T | R473C (CGT→TGT) | *amn* | AMP nucleosidase |

**Supplementary Movie 1 | Time-lapse imaging of wild-type cells during ampicillin treatment.** Time-lapse fluorescence imaging of wild-type cells expressing periplasmic $_{ss}$DsbA–mCherry (red) and cytoplasmic yellow fluorescent protein (YFP; cyan) during 120 µg/ml ampicillin exposure.

**Supplementary Movie 2 | Time-lapse imaging of *amn*$^{T}$ cells during ampicillin treatment.** Time-lapse fluorescence imaging of *amn*$^{T}$ cells expressing periplasmic $_{ss}$DsbA–mCherry (red) and cytoplasmic yellow fluorescent protein (YFP; cyan) during 120 µg/ml ampicillin exposure.

**Supplementary Movie 3 | Time-lapse imaging of wild-type cells during kanamycin treatment.** Time-lapse fluorescence imaging of wild-type cells expressing periplasmic $_{ss}$DsbA–mCherry (red) and cytoplasmic yellow fluorescent protein (YFP; cyan) during 120 µg/ml kanamycin exposure.

**Supplementary Movie 4 | Time-lapse imaging of *amn*$^{T}$ cells during kanamycin treatment.** Time-lapse fluorescence imaging of *amn*$^{T}$ cells expressing periplasmic $_{ss}$DsbA–mCherry (red) and cytoplasmic yellow fluorescent protein (YFP; cyan) during 120 µg/ml kanamycin exposure.

**Supplementary Movie 5 | Time-lapse imaging of *amn*$^{T}$ cells after 10 mM EDTA treatment during ampicillin exposure.** Time-lapse fluorescence imaging of 10 mM EDTA-treated *amn*$^{T}$ cells expressing periplasmic $_{ss}$DsbA–mCherry (red) and cytoplasmic yellow fluorescent protein (YFP; cyan) during 120 µg/ml ampicillin exposure.

**Supplementary Movie 6 | Time-lapse imaging of *amn*$^{T}$ cells after 10 mM EDTA treatment during kanamycin exposure.** Time-lapse fluorescence imaging of 10 mM EDTA-treated *amn*$^{T}$ cells expressing periplasmic $_{ss}$DsbA–mCherry (red) and cytoplasmic yellow fluorescent protein (YFP; cyan) during 120 µg/ml kanamycin exposure.
